# ALLM-Ab: Active Learning-Driven Antibody Optimization Using Fine-tuned Protein Language Models

**DOI:** 10.1101/2025.08.05.668775

**Authors:** Kairi Furui, Masahito Ohue

**Affiliations:** Department of Computer Science, School of Computing, Institute of Science Tokyo, Yokohama 226-8501, Japan

## Abstract

Antibody engineering requires a delicate balance between enhancing binding affinity and maintaining developability properties. In this study, we present ALLM-Ab (Active Learning with Language Models for Antibodies), a novel active learning framework that leverages fine-tuned protein language models to accelerate antibody sequence optimization. By employing parameter-efficient fine-tuning via low-rank adaptation, coupled with a learning-to-rank strategy, ALLM-Ab accurately assesses mutant fitness while efficiently generating candidate sequences through direct sampling from the model’s probability distribution. Furthermore, by integrating a multi-objective optimization scheme incorporating antibody developability metrics, the framework ensures that optimized sequences retain therapeutic antibody-like properties alongside improved binding affinity. We validate ALLM-Ab in both offline experiments using deep mutational scanning (DMS) data from the BindingGYM dataset and online active learning trials targeting Flex ddG energy minimization across three antigens. Results demonstrate that ALLM-Ab not only expedites the discovery of high-affinity variants compared to baseline Gaussian process regression and genetic algorithm-based approaches, but also preserves critical antibody developability metrics. This work lays the foundation for more efficient and reliable antibody design strategies, with the potential to significantly reduce therapeutic development costs.

## Introduction

Optimizing the binding affinity between proteins is a critical aspect of drug development, including antibody engineering.^1,2^ In recent years, deep learning-based *de novo* antibody design methods have been proposed.^3–8^ However, approaches that leverage existing antibody information to design improved variants are also important, as *de novo* methods may not fully exploit the prior knowledge of existing antibodies. This is particularly relevant when dealing with known antibodies that require improvements in desirable properties such as binding affinity, specificity, and developability.

Large-scale protein language models (pLMs)^9–12^ inspired by natural language processing^1^^3^ treat amino acid sequences as a “language” and learn the statistical patterns of protein sequences from vast databases, achieving state-of-the-art performance in protein structure and function prediction.^10,11^ In particular, pLMs can evaluate the fitness of mutant sequences, which often correlates with their functional fitness, enabling the prediction of beneficial mutations without additional training.^11,14^ However, because basic pLMs are not specifically trained for design objectives like binding affinity to target proteins, their accuracy for predicting such specific properties may be limited.^15^ Few-shot learning approaches that fine-tune language models with limited data have shown promising results.^15,16^

In recent years, active learning^17–19^—a machine learning approach—has attracted attention in drug development as a means to streamline computationally expensive experiments such as compound docking and molecular simulations. ^20–22^ Active learning uses a surrogate model to iteratively select the next data points to evaluate.^17–19^ This approach is designed to efficiently collect data by balancing an exploration phase, which searches regions of high uncertainty, with an exploitation phase that seeks data with desirable properties. Even with limited experimental data, active learning can efficiently explore promising candidates by iteratively adding new data, thus advancing optimization more effectively than traditional exhaustive screening methods.

In the field of small molecule drug development, active learning combined with molecular simulations and docking simulations has successfully enhanced screening efficiency without relying on experimental data.^20–22^ Similarly, in antibody engineering several active learning approaches have been proposed.^23–27^ Several approaches have proposed methods for optimizing sequences using Bayesian optimization^27^ or deep learning techniques such as Transformers^23^ and Variational Autoencoders,^26^ trained on datasets with thousands or more sufficient training examples. However, these methods assume the availability of thousands of training examples and are difficult to apply in situations where little training data exists. On the other hand, some approaches have proposed methods for constructing active learning models from a small number of sequence data,^24,25,28^ demonstrating that promising variants can be obtained even when little known experimental data is available. Khan *et al.*^24^ proposed a combinatorial Bayesian optimization tool called AntBO. Furthermore, Gessner *et al.*^25^ successfully discovered antibody sequences with improved binding affinity based on relative binding free energies computed using Bayesian optimization.

Despite advances in computational methods for antibody active learning with limited sequence data, several challenges remain. First, it is possible that the extensive prior knowledge embedded in prevalent language models has not been fully exploited. For example, approaches such as Gaussian process regression (GPR) that use the latent space of a language model as features^24,25^ may lose detailed residue and mutation-specific information, thereby failing to fully integrate the vast prior knowledge from protein language models with the experimental data. As a result, surrogate models based solely on limited experimental data may not capture the complex patterns of antibody-antigen interactions, potentially limiting performance. Indeed, in Khan *et al.*’s experiments,^24^ the performance of a Gaussian process model using kernels derived from ProteinBERT,^29^ a protein language model, was inferior to simpler methods such as kernels based on sequence Hamming distance.

Moreover, although fine-tuning language models has shown promising results in predicting the fitness of single mutations,^15^ how to efficiently generate mutant sequences from the fine-tuned models remains an open challenge. For example, traditional approaches using genetic algorithms (GA)^25,30^ rely on random mutation operations, which do not directly leverage the prior knowledge acquired by the language models, making it difficult to efficiently explore the vast sequence space.

Furthermore, when improving antibody binding affinity, it is necessary to simultaneously avoid undesirable properties for practical drug development, such as non-specificity, self-association, and instability. ^31–33^ Optimizing these properties in a comprehensive manner requires a multi-objective optimization approach.

To address these challenges, we propose ALLM-Ab (Active Learning with Language Models for Antibodies), a comprehensive active learning framework that leverages fine-tuned protein language models. In our approach, both the inference and generation processes utilize the prior knowledge of the language models as well as the knowledge acquired from limited data, thereby efficiently optimizing binding affinity while preserving the natural characteristics of antibodies.

Our approach consists of three components. First, we construct a surrogate model for active learning using a protein language model by employing parameter-efficient fine-tuning(PEFT)^34,35^ and learning-to-rank^36–38^ as proposed by Zhou *et al.*^15^ By leveraging the general knowledge of the pLM, the surrogate model can learn the specific interaction patterns between antibodies and antigens even from limited training data, enabling a more accurate search for high-affinity mutants. Next, the fine-tuned pLM is integrated into an active learning pipeline that not only evaluates mutant sequences but also generates new candidate sequences. Instead of selecting from a predetermined library, candidate sequences are directly sampled from the probability distribution of the fine-tuned pLM, thereby improving exploration efficiency. Finally, we incorporate multi-objective active learning based on hypervolume maximization. When sampling sequences using binding information-based fine-tuning, there is a risk of over-optimization toward high affinity scores, which may result in the generation of sequences with low validity as antibodies. To counteract this, we incorporate developability metrics such as the perplexity score from AbLang2, an antibody language model pre-trained on the OAS sequence database,^39,40^ as well as hydropathicity, ^41^ instability,^42^ and isoelectric point^43^ calculated from sequences as constraints. This constraint prevents sequences selected from the overfitted fine-tuned model from straying too far from the original antibody sequence space, thereby enabling the discovery of sequences with high fitness while preserving valid sequence patterns as therapeutic antibodies.

To evaluate the effectiveness of the proposed approach, we conducted experiments in two scenarios: (i) offline active learning using a DMS benchmark dataset and (ii) online active learning that includes mutant generation and is aimed at improving Flex ddG’s energy values. In the offline setting, we simulated the active learning process using deep mutational scanning (DMS) data from the BindingGYM dataset. In this experiment, no new sequences were generated; instead, mutants were selected from a pool with known binding scores to evaluate whether ALLM-Ab can efficiently identify high-affinity mutants. In the online setting, we performed an active learning trial using a Rosetta-based computational method called Flex ddG. Here, the fine-tuned model directly generates mutants, and we evaluated whether this approach can explore mutants with higher affinity compared to generating sequences in advance. Moreover, by performing multi-objective optimization that incorporates both the binding score and the perplexity from AbLang2^40^ along with developability metrics, we investigated whether it is possible to prevent excessive optimization toward Flex ddG scores. Finally, we conducted experiments on bispecific antibodies targeting two antigens, 5A12_Ang2 and 5A12_VEGF, to assess whether ALLM-Ab can generate more practical antibody sequences through multi-objective active learning. The results of these experiments demonstrate that the proposed approach can efficiently optimize antibody binding affinity while preserving the natural sequence features of antibodies.

## Materials and Methods

In this section, we describe the core components of ALLM-Ab, our proposed active learning framework for antibody optimization and the details of each component. Then, we provide detailed explanations of both the offline experimental setup using the BindingGYM dataset and the setup for Flex ddG’s energy Improvement via Online Active Learning.

### Multi-Objective Batch Online Active Learning

An overview of the proposed method, ALLM-Ab, is shown in Figure 1. Our active learning framework is composed of three components:

1. **Parameter-efficient fine-tuning and learning-to-rank of the pLM:** We perform few-shot learning of the protein language model using PEFT^34,35^ and learning-to-rank^36–38^ as proposed by Zhou *et al.*^15^
2. **Mutant generation using the fine-tuned model:** We generate candidate sequences by sampling from the probability distribution of the fine-tuned model. By directly sampling mutants that reflect the model’s adjusted preferences, ALLM-Ab improves the efficiency of online active learning. Furthermore, we accelerate inference by utilizing an approximate fitness score.
3. **Multi-objective active learning based on hypervolume maximization:** When sampling sequences using binding information-based fine-tuning, there is a risk of over-optimization toward high scores, which may result in the generation of sequences with low validity as antibodies. ^44^ To address this challenge, we incorporate the binding score, the perplexity score from AbLang2, and developability metrics such as hydropathy, instability, and appropriate isoelectric point, aiming to select antibodies that are both natural and have high developability. We perform sequence selection based on hypervolume maximization^45^ to optimize these multiple objectives simultaneously.

**Figure 1:**
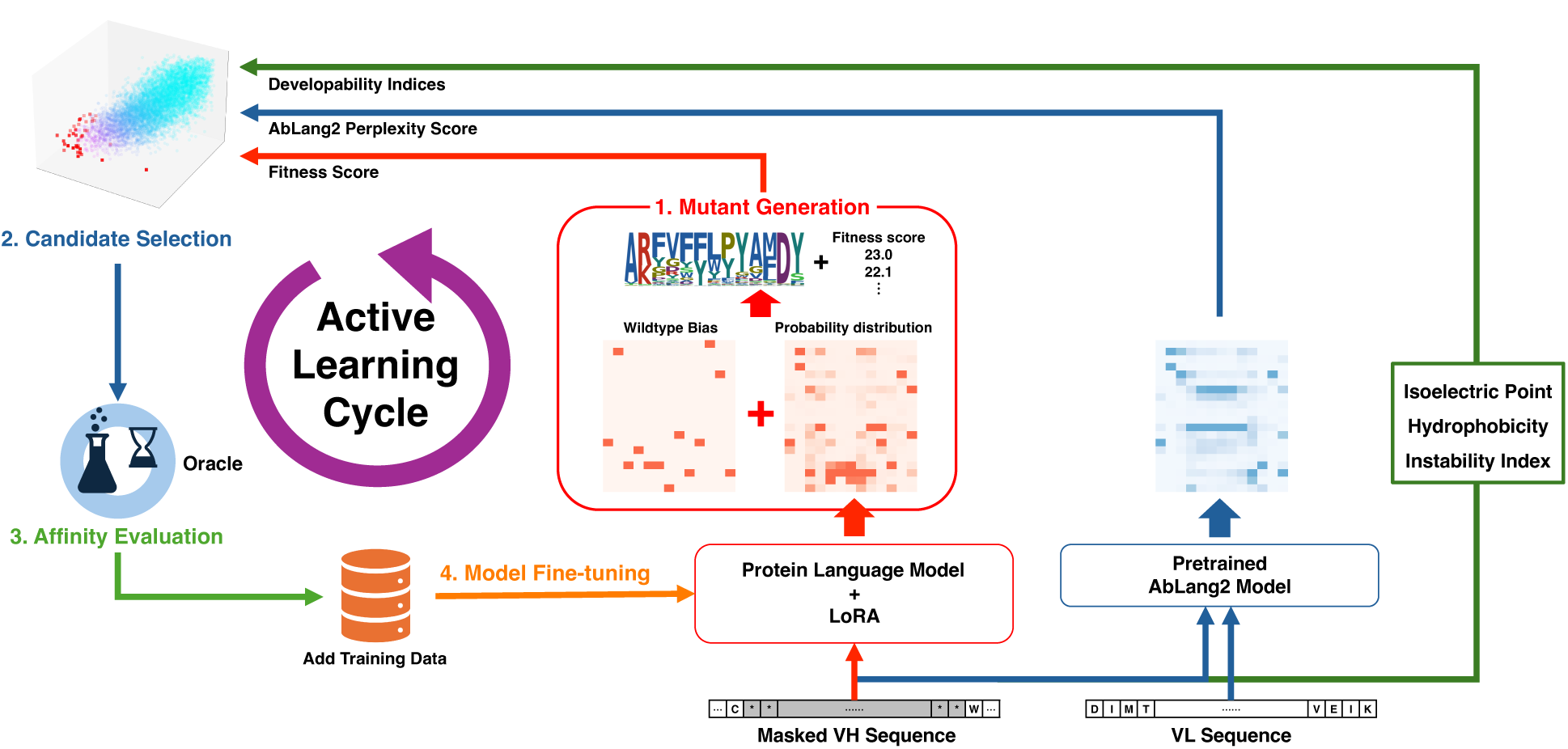
Overview of the proposed method, ALLM-Ab.

Based on the aforementioned components, the active learning workflow proceeds as follows. First, sampling is performed from the probability distribution of a fine-tuned protein language model (or a pre-trained protein language model in the initial cycle). Next, multi-objective optimization is conducted based on fitness scores for the sampled sequences, AbLang2 perplexity, and developability metrics. Selected sequences are then evaluated using an oracle optimization target such as biochemical experiments or energy calculations to obtain affinity scores. Finally, the protein language model is fine-tuned based on the acquired affinity scores, and the procedure is repeated.

### Fitness Score Using pLMs

In active learning with protein language models, the difference in log-likelihood between the wild-type sequence and its mutant is used as the fitness score. Specifically, following previous work,^46^ the fitness score *f* is defined as:

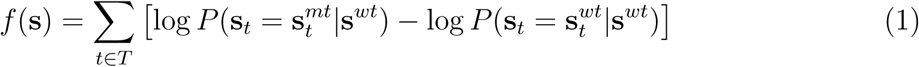

Here, 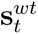 and 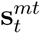 denote the *t*-th amino acid of the wild-type and mutant sequences, respectively, and *T* represents the set of all mutated positions in **s***^mt^*. This score is computed in the same way for both zero-shot and fine-tuned models.

Furthermore, as in Zhou *et al.*,^15^ protein language models such as ESM-2^9^ and AbLang2^40^ are fine-tuned via low-rank adaptation,^35^ which modifies only a subset of parameters. This approach allows the prediction of target-specific fitness from limited data while preventing overfitting, since only a very small number of parameters are adjusted compared to fine-tuning the entire pLM.

Moreover, the model is fine-tuned using a learning-to-rank based on ListMLE.^36,37^ In this loss function, the fitness scores predicted by the protein language model are optimized to accurately rank the mutant sequences. In active learning, it is more important that the mutants ranked highest by the model correspond to high actual binding affinity than to accurately predict the exact binding affinity values. This learning-to-rank approach better aligns the fitness function with active learning goals, leading to more efficient mutant exploration. Because computing the fitness score *f* requires inferring the masked probability for only the mutated regions of each mutant sequence, evaluating a large number of sequences can be computationally expensive. To improve inference speed, we pre-compute the probability distribution for a masked sequence **s***^mask^*—in which all potentially mutable regions (e.g., the entire CDR-H3) are masked—for each task, and use this distribution to approximate the fitness score *f*. We define this modified score, called the approximation score *f_approx_*, as follows:

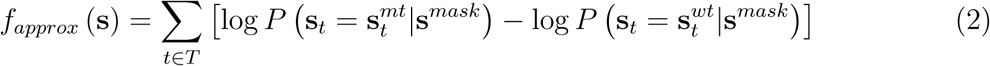

In this formulation, the probability distribution *P* (**s***_t_*| **s***^mask^*) is computed only once for **s***^mask^*, enabling rapid evaluation of the score for multiple mutant sequences.

### Mutant Generation Using the Fine-tuned Model

In active learning using pLMs, since amino acid sequences are discrete variables, it is difficult to directly apply continuous optimization methods^47^ that directly explore the feature space using Bayesian optimization.^48,49^ Additionally, exploration methods such as genetic algorithms (GA)^25,30^ face challenges in exploration efficiency in vast sequence spaces.

Therefore, we propose to generate sequences by sampling from the fine-tuned model’s probability distribution, which directly incorporates the model’s preferences to improve active learning efficiency. Even masked language models such as ESM2 can generate sequences from the probability distribution of a masked sequence. ^12,50,51^ Specifically, for all mutable amino acid positions in the set *T*, we pre-compute the probability distribution of the masked sequence **s***^mask^* and then perform sampling at each residue position according to this distribution. This **s***^mask^* can also be used for calculating the approximation score.

Here, we sample the residue **s***_t_* at position *t* with probability *P* ^′^ as follows using the logits computed by the pLM from **s***^mask^*:

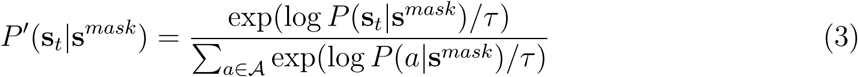

where 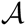 denotes the set of the standard 20 amino acids and *τ* is a temperature parameter. The higher the temperature parameter *τ*, the more likely amino acids with lower probabilities are to be sampled. This temperature parameter is used to adjust the balance between exploration and exploitation.

Furthermore, to preserve the characteristics of the wild-type sequence, a correction is applied using the wild-type amino acids **s***^wt^* as follows, and sampling is performed with probability *P_biased_*:

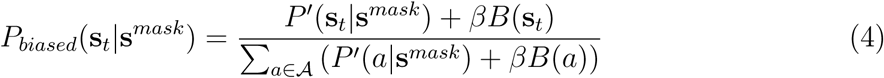

Here, 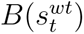 is a bias probability that assigns a probability of 1 only to the amino acid present at the *t*-th position of the wild-type sequence, and *β* is a parameter representing the strength of the bias. The larger *β* is, the more the sequences are sampled with emphasis on wild-type residues. The bias toward wild-type sequences was introduced with the aim of preventing excessive deviation from known sequence spaces by exploring sequences based on wild-type sequences as a reference.

Note that any generated mutants containing a solitary cysteine residue are filtered out.

This approach enables sampling that reflects the preferences of the fine-tuned model while keeping the generated sequences close to the wild-type sequence.

### Multi-Objective Active Learning

In this study, we perform multi-objective active learning using the binding score, the perplexity score from the pre-trained AbLang2, ^40^ and several developability metrics calculated from sequences as objective functions, with the goal of discovering sequences that have high fitness while maintaining their validity as antibody sequences.

AbLang2^40^ is pre-trained on paired heavy and light chain sequences from the OAS database^39^ and implicitly learns how valid a sequence is as an antibody. Likelihood scores from protein language models have been shown to predict missense mutations and stability in zero-shot.^46,52^ Furthermore, log-likelihood scores derived from antibody language models like AbLang2 have been demonstrated to correlate with experimental binding affinities without antigen information.^53^ Therefore, by optimizing the perplexity score of antibody language models, we can implicitly incorporate antibody-specific preferences into the design criteria. In this work, we evaluate the validity of an antibody sequence by defining the AbLang2’s perplexity (hereafter AbLang2 perplexity score) as follows:

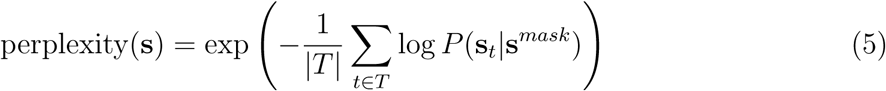

where *T* denotes the set of mutated positions. The score is calculated as the exponential of the negative mean log-likelihood over the mutated positions, where lower values indicate greater consistency with natural antibody sequence patterns.

Additionally, we aim to discover sequences with high binding affinity while maintaining desirable sequence patterns by incorporating the following three developability metrics as objective variables:

1. **Isoelectric point**: The isoelectric point is the pH value at which the net charge of a protein is zero.^43^ This is to avoid the risks of non-specific binding and self-association from excessively high or low IP values. ^31^ Based on previous research^54^ observing that antibodies with isoelectric points in the range of 6.7 to 9.05 have slow clearance, the objective function was designed so that the isoelectric point would be 8.
2. **Hydropathicity**: High hydropathicity promotes non-specific binding, so this is avoided. It was calculated based on the GRAVY (Grand average of hydropathicity) score. ^41^
3. **Instability index**: To evaluate the stability of designed sequences, the instability index^42^ should not become large. The instability index is considered stable if it is less than 40.

These metrics were calculated based on the VH region sequence using Biopython.

Furthermore, for multi-objective active learning, we select multiple mutants based on improvements in the hypervolume,^45^ which enables the selection of mutants that simultaneously optimize the scores of multiple objective functions. First, for each objective function score *S* of the generated mutants, we normalize it using the 5th percentile *S*_5%_ and the 95^th^percentile *S*_95%_:

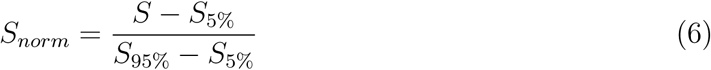

Here, the fitness score is multiplied by −1 to transform it into a quantity to be minimized. Using these normalized scores, we select a set of mutants that maximizes the hypervolume *HV*.

In a multi-objective optimization with *M* objective functions, let the Pareto solution set be 𝒫 = {*x*_1_*, x*_2_*,…, x_n_*}, and let the objective vector of each solution *x_i_* be **S**(*x_i_*) = (*W*_1_*S*_1_(*x_i_*)*, W*_2_*S*_2_(*x_i_*)*,…, W_M_ S_M_* (*x_i_*)). Here, *W_j_* for each objective variable *j* represents the weight for each objective variable, and the larger it is, the more it contributes to the hypervolume calculation, thus expressing importance. When the reference point is **r** = (*r*_1_*, r*_2_*,…, r_M_*), the hypervolume *HV* (𝒫) is defined as follows:

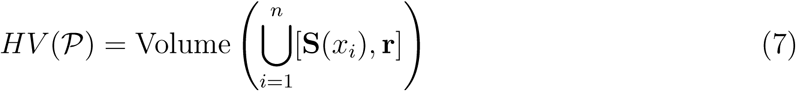

where [**S**(*x_i_*), **r**] represents a hypercube with solution **S**(*x_i_*) and reference point **r** as diagonals.

The set of *n* mutants that maximizes the hypervolume is selected in a greedy manner as follows. First, the reference point is set to the 95th percentile (*S*_95%_) of each objective function. Any mutant whose score in any objective exceeds the 95th percentile is excluded from the candidate set, as it cannot be used in the hypervolume calculation. Let *V* denote the set of already selected mutants and *R* the set of remaining candidates. At each step, for every mutant in *R*, we compute the hypervolume resulting from adding that mutant to *V*, and we select the mutant that yields the maximum increase in hypervolume. If no further improvement in hypervolume is observed, the remaining mutants are selected in descending order of their fitness score. The hypervolume is computed using the combined set of selected mutants *V* and the candidate mutant. By repeating the operation of adding the chosen mutant to *V* and removing it from *R* for *n* iterations, a set of *n* mutants that maximizes the hypervolume is obtained. The computational complexity for evaluating the hypervolume when selecting *n* mutants from *N* is *O*(*nN*). The hypervolume calculation is performed using pygmo.^55^

Finally, mutants are ranked in ascending order of promising mutants by obtaining a weighted sum of the obtained normalized scores.

## Experiments

### Offline Active Learning Evaluation on the BindingGYM Dataset

In this experiment, we evaluate ALLM-Ab using three target datasets selected from the DMS data of the BindingGYM dataset.^56^ BindingGYM is a deep mutational scanning (DMS) dataset focused on protein-protein interactions, containing 10 million data points. In this experiment, we explore only a pool of known DMS dataset, aiming to identify mutants with higher DMS scores rather than generating new sequences (see Figure 2). For our evaluation, we use three antigen-antibody complex datasets related to therapeutic antibody design. The details of each dataset are shown in Table 1. Among these, the 5A12 antibody targeting 5A12_Ang2 and 5A12_VEGF is bispecific,^57,58^ and the wild-type sequence is identical.

**Figure 2:**
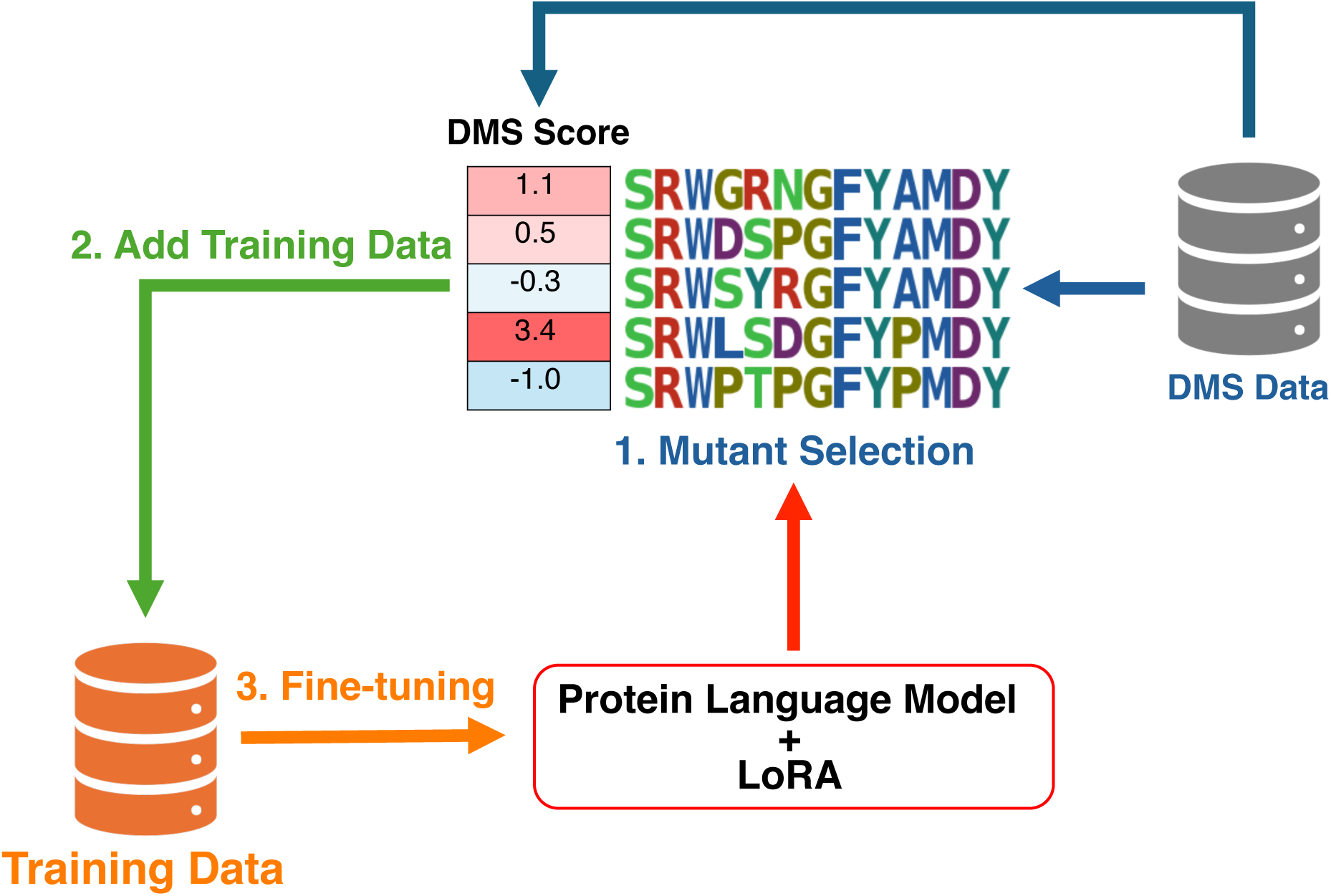
Overview of offline active learning.

**Table 1:** Details of the BindingGYM datasets used in offline active learning.

| Label | Protein 1 | Protein 2 | #Variants | PDBID |
| --- | --- | --- | --- | --- |
| 4D5_HER2<sup>58,59</sup> | 4D5 | HER2 | 2,080 | 1N8Z<sup>60</sup> |
| 5A12_Ang2<sup>58</sup> | 5A12 | Ang2 | 944 | 4ZFG<sup>57</sup> |
| 5A12_VEGF<sup>58</sup> | 5A12 | VEGF | 29,981 | 4ZFF<sup>57</sup> |

From these data, 100 samples are randomly selected as a test set, and the predictive performance of the active learning models is evaluated using the Spearman correlation on the test set. For each target, offline active learning is performed with the aim of identifying mutants with higher DMS scores.

We set the number of mutants selected per cycle as *N_cycle_* = 50, and select a total of 600 mutants over multiple cycles.

#### Model Training

As ALLM-Ab, we compare four models: two methods based on fine-tuning of language models (ESM2^9^ and AbLang2^40^), a method based on fine-tuning of the structure-based inverse folding model ProteinMPNN,^61^ and an existing method using Gaussian process regression (GPR) based on the latent variables of AbLang2 (denoted as GPR(AbLang2)).

For ESM2, we use `esm2_t33_650M_UR50D`. Following Zhou *et al.*,^15^ we fix all parameters except those in the attention layers and some linear layers, and fine-tune using LoRA (Low-Rank Adaptation).^35^ LoRA rank was set to 8 and dropout rate to 0.1, with all other parameters using default values. For AbLang2, we use paired variable region sequences and employ LoRA for fine-tuning. The specific layers to which LoRA was applied are listed in Table S1. For ESM2, when mutations span multiple chains, sequences are concatenated as described in Lu *et al.*^56^ Unless otherwise stated, the fitness score *f* is used for inference. For ProteinMPNN, we fine-tune the pre-trained full-protein backbone model `v_48_020`. For ProteinMPNN, fitness scores are calculated following the method of Lu *et al.*^56^ Unlike their approach of using transfer learning with the entire BindingGYM dataset, we incorporated LoRA into ProteinMPNN to prevent overfitting due to our limited training data. For ESM2, AbLang2, and ProteinMPNN, we use the ListMLE loss^37^ as in Zhou *et al.*,^15^ emphasizing the accurate prediction of the ranking order of mutant sequences.

To evaluate the model’s performance, datasets containing more than 100 samples are split into training and validation sets in a 4:1 ratio. For datasets with fewer than 100 samples, we determine the optimal number of epochs via Monte Carlo cross-validation following the method of Zhou *et al.*^15^ using 5-fold cross-validation. The optimal number of epochs is then used to retrain the model on the entire dataset, which helps to avoid overfitting when data are limited.

As a baseline, we also employ Gaussian process regression (GPR) using the 480-dimensional latent variables from AbLang2. ^25^ In GPR(AbLang2), the model learns the actual DMS score values rather than their ranking. The kernel function is a combination of a constant term, white noise, and a radial basis function (RBF).

#### Evaluation Metrics

In this study, active learning performance is evaluated using the metrics TopMean@10, Top 1% Recall, and Spearman’s *ρ*. Specifically:

**TopMean@10** The average DMS score of the top 10 mutants selected by active learning.

A higher value indicates that more high-scoring mutants were discovered.

**Top 1% Recall** The proportion of positive mutants (defined as the top 1% based on DMS scores) that are included in the active learning selection. A higher recall indicates that more promising mutants were identified.

**Spearman’s** *ρ* The Spearman’s rank correlation coefficient between the experimental DMS scores and the model’s predicted fitness scores on the test set.

### Flex ddG’s Energy Improvement via Online Active Learning

The offline active learning experiments using known DMS data are limited to existing mutants and do not address the exploration of unknown mutants. Therefore, we conducted online active learning validation using 15 targets by adding 12 targets used in AntBO experiments^24^ to the 3 targets from the offline experiments, exploring mutant sequences with only the wild-type sequence known. In this setting, we aim to improve the ΔΔ*G_FlexddG_* values related to protein-protein binding mutations as calculated by the Rosetta-based method Flex ddG.^56^ Although Flex ddG achieves performance comparable to state-of-the-art methods,^62^ its high computational cost is a known challenge. We evaluate the performance of ALLM-Ab by exploring mutants that optimize the Flex ddG values. It should be noted that this experiment uses computational methods rather than actual biochemical experiments, and is intended as a proof-of-concept to validate the performance of ALLM-Ab in a controlled environment where ground truth values are available.

**Table 2:** Details of targets in online active learning.

| Source | PDB | Target |
| --- | --- | --- |
| BindingGYM | 1N8Z_C | HER2: Receptor protein-tyrosine kinase erbB-2 (Trastuzumab) |
|  | 4ZFG_A | Ang2: Angiopoietin 2 |
|  | 4ZFF_C | VEGF: vascular endothelial growth factor |
| AntBO | 1ADQ_A | IGG4 Fc Region |
|  | 1FBI_X | Guinea Fowl Lysozyme |
|  | 1H0D_C | Angiogenin |
|  | 1NSN_S | SNASE: Staphylococcal nuclease complex |
|  | 1OB1_C | MSP1: Merozoite Surface Protein 1 |
|  | 1WEJ_F | CYC: Cytochrome C |
|  | 2YPV_A | fHbp: factor H binding protein |
|  | 3RAJ_A | CD38: ADP-Ribosyl cyclase 1 |
|  | 3VRL_C | HIV Gag protein |
|  | 2DD8_S | SARS-CoV Virus Spike glycoprotein |
|  | 1S78_B | HER2: Receptor protein-tyrosine kinase erbB-2 (Pertuzumab) |
|  | 2JEL_P | ptsH: Phosphocarrier protein HPr |

#### Experimental Setting

In our batch active learning framework, *N_cycle_* = 40 mutants are selected per cycle for a total of 10 cycles (i.e., 400 mutants), and the Flex ddG energy values of these mutants are evaluated. Starting from the wild-type sequence, we focus only on the CDR H3 region of the antibody and search for mutants that lower the Flex ddG energy values. During inference, the approximation score *f_approx_* is used to accelerate the process. To reduce computational cost, only a single optimized energy value is computed per mutant without employing ensemble structures in Flex ddG; all other parameters follow the default settings of Flex ddG.

Three experimental settings are considered:

##### Single-Objective Sequence Sampling

Active learning is performed using only the Flex ddG energy as the objective function. The performance of different sequence generation methods is compared based on their ability to discover sequences with low energy values.

##### Multi-Objective Optimization

Active learning is performed by adding multiple antibody developability metrics as objective function in addition to the Flex ddG energy. The antibody developability and binding affinity are evaluated for the use of different objective functions.

##### Dual Optimization for Ang2 and VEGF

Active learning is conducted for bispecific antibodies targeting both 5A12_Ang2 and 5A12_VEGF using the two objective functions (Flex ddG energy and developability metrics). Both ALLM-Ab and FlexddG-Only settings are evaluated.

For the sequence sampling based on the fine-tuned language models, we compare the following five methods:

##### Online (biased)

At each active learning cycle, mutants are generated by sampling from the probability distribution of the fine-tuned model using Eq. 4, which applies bias from the wild-type sequence.

##### Online (unbiased)

Mutants are generated by sampling from the fine-tuned model using Eq. 3 without applying any bias.

##### Offline (biased)

Candidate mutants are generated using the pre-trained model with bias applied as in Eq. 4.

##### Offline (unbiased)

Candidate mutants are generated using the pre-trained model without bias as in Eq. 3.

##### GA

A genetic algorithm-based sequence selection method is used, as described in the next section.

Here, the bias toward the wild-type sequence was set to *β* = 1, and the temperature parameter was set to *τ* = 1.

For online-based methods, 10,000 candidate mutants are generated in each cycle, from which *N_cycle_* = 40 are selected. For offline-based methods, 100,000 mutants are pre-generated using the pre-trained model before starting active learning. The GA method is executed independently for *N_cycle_* iterations, selecting one mutant from each independent run to maintain diversity in batch active learning. For GPR(AbLang2), only the two offline methods and the GA-based method were evaluated. For dual antigen optimization, the probability distributions of two fine-tuned models are summed to generate candidate sequences.

We evaluate the performance of AbLang2 and GPR(AbLang2) models in this experiment. In addition, a test set of 400 sequences generated using the pre-trained AbLang2 is used to evaluate the predictive performance by computing their Flex ddG values.

To evaluate the multi-objective optimization performance of ALLM-Ab, we compared the following five methods:

##### FlexddG-Only

Optimize with only the fitness score as the objective variable.

##### ALLM-Ab+A

Multi-objective optimization with two objective variables: fitness score and AbLang2 perplexity score.

##### ALLM-Ab+AP

Add isoelectric **P**oint to ALLM-Ab+A.

##### ALLM-Ab+APH

Add **H**ydropathy to ALLM-Ab+AP.

##### ALLM-Ab+APHI

Add **I**nstability index to ALLM-Ab+APH. Hereafter, this is simply referred to as ALLM-Ab.

##### AntBO

Comparison with existing AntBO.^24^

For multi-objective optimization, the fitness score was weighted at 2.0, while other metrics were weighted at 1.0 when included in the optimization and 0 when excluded, thereby prioritizing fitness score optimization. During performance evaluation, the top 40 cases are evaluated in ascending order of the weighted sum of normalized scores in the final cycle.

#### Sequence Generation via Genetic Algorithms

For comparison with ALLM-Ab, a genetic algorithm (GA)-based sequence generation method was employed. In the GA-based approach, the initial population includes the wild-type sequence, and the remaining sequences are generated by randomly selecting amino acids. Genetic operations include two-point crossover and uniform mutation. Elitist selection is used, where the highest-fitness individual is carried over unchanged to the next generation. The parameters are set as follows: population size of 30, 10 offspring generated per generation, and 100 generations; crossover probability is 0.7, and mutation probability is 0.1. In the GA-based sequence generation, the DEAP framework^63^ is used.

It was observed that the population generated by the GA tends to exhibit low diversity due to optimization based on a single objective function. In batch active learning, this reduction in diversity can be problematic for efficiently exploring the search space. As a naive solution, we executed the GA independently for *N_cycle_* iterations, selecting one mutant from each independent run to maintain diversity.

### Comparison with Existing Methods

As a competing existing method, we compare with a method called AntBO.^24^ AntBO is a combinatorial Bayesian optimization framework for antibody CDR-H3 region design that sequentially generates new candidate sequences by optimizing acquisition functions within a trust region that satisfies developability constraints. In AntBO, three conditions are set as developability constraints: the net charge of the sequence is within the range of −2 to +2, consecutive occurrences of the same residue are 5 times or less, and no glycosylation motifs are included. This enables antibody design that considers practical development requirements such as developability and stability. In this study, we use the transformed overlap kernel, which performed best in their research, as the kernel for AntBO. Then, giving only the wildtype CDR-H3 as the initial sequence, we search for sequences that improve the Flex ddG energy value with a batch size of 40. We use EI as the acquisition function for AntBO and set other parameters based on the values in their paper.

#### Evaluation Metrics

In single-objective optimization settings, the performance of surrogate models is evaluated by tracking the progression of TopMean@40 and Spearman correlation on the test set. TopMean@40 is defined as the average Flex ddG value of the top 40 mutants selected up to that cycle.

In multi-objective optimization and dual optimization experiments, mutants are first ranked by the weighted sum of normalized objective scores, and each objective metric is evaluated for the top 40 mutants. Furthermore, to evaluate antibody developability using external criteria, we assess 8 developability metrics for the top-performing mutants from each method per target. These metrics include PSH, PPC, PNC, SFvCSP from therapeutic antibody profiler (TAP),^64^ Heavy OASis Percentile from BioPhi,^65^ and `SAP_pos_CDR-H3, SCM_pos_CDR-H3`, `SCM_neg_CDR-H3` from DeepSP.^66^ TAP is a tool that generates antibody model structures from antibody variable domain sequences using ABodyBuilder2^67^ and checks whether they match the characteristics of clinical-stage therapeutics. BioPhi is an antibody design platform for evaluating the human-likeness of antibodies. DeepSP is a convolutional neural network-based deep learning model that predicts spatial properties related to antibody stability from antibody sequence information alone.

The meanings of each metric are as follows:

##### PSH

Patches of Surface Hydrophobicity. Must be within acceptable range. An indicator of the degree of surface hydrophobic patches near CDRs, where higher values indicate the presence of hydrophobic residues in close proximity. High values suggest risks of non-specific binding or high blood clearance.^64^

##### PPC

Patches of Positive Charges. ^68^ When too high, there are risks of non-specific binding or high blood clearance, and it needs to be less than 0.73.

##### PNC

Patches of Negative Charges. ^69^ When too high, it can cause decreased expression levels or structural instability issues, and it needs to be less than 1.28.

##### SFvCSP

Structural Fv charge symmetry represents the asymmetry of surface charges between heavy and light chains.^70,71^ Larger negative values increase the risk of viscosity increase, and it needs to be −1.59 or higher.

##### Heavy OASis Percentile

Heavy chain humanization score based on 9-mer peptide search in the Observed Antibody Space (OAS) calculated by BioPhi, where larger values indicate greater human-likeness.

##### SAPpos

Spatial aggregation propensity of CDR-H3 calculated by DeepSP, where larger values indicate more hydrophobic regions and higher aggregation risk.

##### SCMpos

Positive Spatial Charge Map of CDR-H3 calculated by DeepSP, where extremely high values pose aggregation risks.

##### SCMneg

Negative Spatial Charge Map of CDR-H3 calculated by DeepSP.

These metrics were calculated using Tamarind.^72^ Note that TAP uses values and thresholds obtained from Tamarind’s reproduced implementation, which differ from the original by Raybould *et al.*^64^

## Results and Discussion

### Offline Active Learning Evaluation

First, we present the results of offline active learning. Figure 3 shows the evolution of active learning performance in terms of TopMean@10, Top 1% Recall, and Spearman’s *ρ* metrics. Figure 4 displays scatter plots comparing the experimental DMS scores and the model predictions on the test set. Overall, models achieving higher Spearman correlation on the test set tended to show better active learning performance in terms of both TopMean@10 and Top 1% Recall metrics. For all targets, GPR(AbLang2) achieved higher active learning performance on average, with a particularly notable improvement in Top 1% Recall for 5A12_VEGF. For 4D5_HER2, model performance was relatively similar; for the other targets, GPR(AbLang2) generally performed best. However, for 5A12_Ang2, both the Spearman correlation and the active learning performance did not improve much over the zero-shot model.

**Figure 3:**
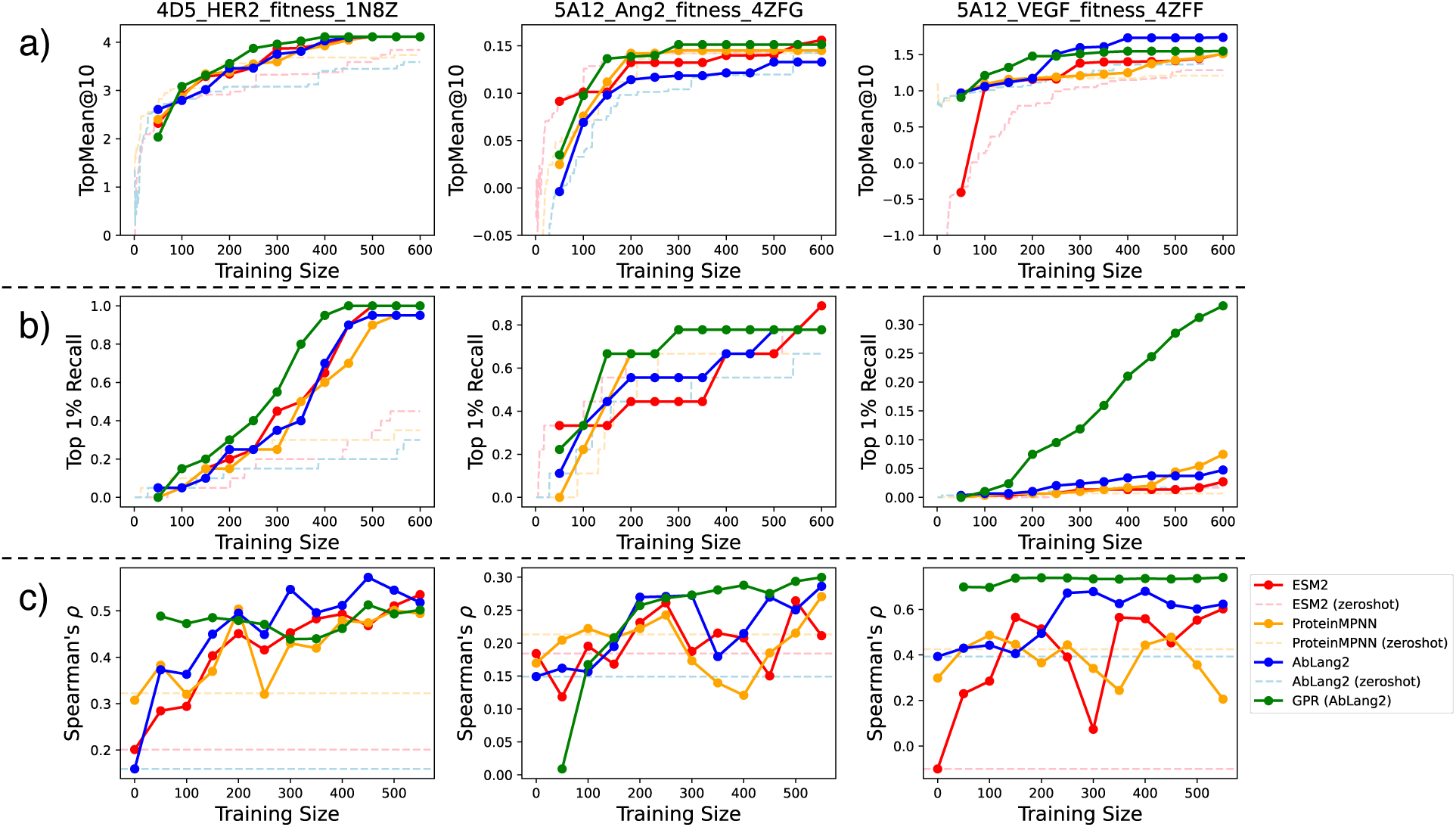
Evolution of active learning performance for each target and model. (a) TopMean@10, (b) Top 1% Recall, (c) Spearman’s *ρ*.

**Figure 4:**
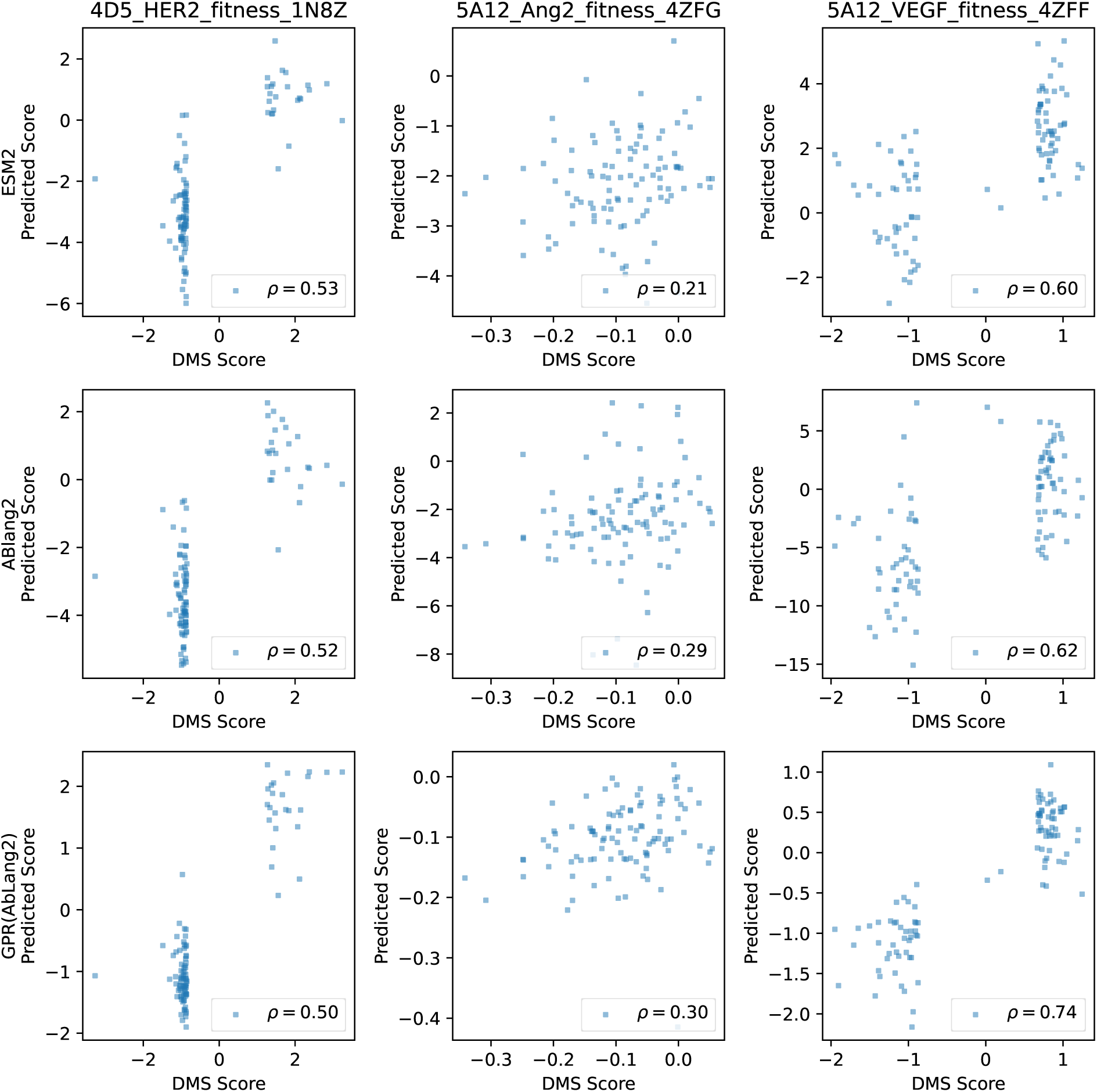
Scatter plots of DMS scores versus predicted scores on the test set for each target and model.

While ProteinMPNN showed the highest predictive performance overall in the zero-shot setting, after fine-tuning there was no clear advantage of ProteinMPNN over the language model-based methods. This suggests that although ProteinMPNN may perform better in the zero-shot setting or with ample training data (as shown in Lu *et al.*^56^), language models are better suited for efficient fine-tuning with few samples.

Based on these results, when performing offline active learning using pre-existing DMS data, employing GPR(AbLang2) appears to yield higher performance. However, since this experiment assumes the existence of mutant data, evaluating active learning performance in the context of mutant generation (as in the next section) is even more critical.

Next, Figure 5 compares the active learning performance when using the direct fitness score (normal mode) versus the approximation score (approx mode) during inference. For ESM2, the Spearman correlation in approx mode was lower than in normal mode, whereas for AbLang2 the performance difference was negligible. In terms of TopMean@10, only the normal mode of AbLang2 was slightly higher; however, in Top 1% Recall both ESM2 and AbLang2 achieved higher performance in approx mode. These results indicate that, for active learning purposes, using the approximation score can dramatically reduce inference time without substantially degrading active learning performance.

**Figure 5:**
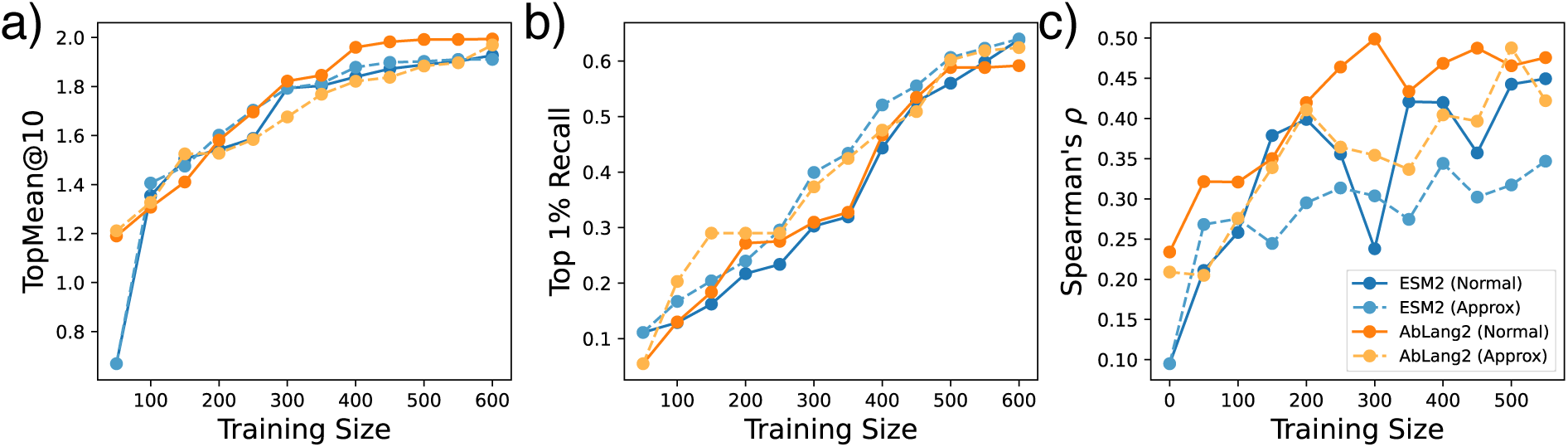
Comparison of active learning performance when using the fitness score (normal mode) versus the approximation score (approx mode). (a) TopMean@10, (b) Top 1% Recall, (c) Spearman’s *ρ*.

### Flex ddG’s Energy Improvement via Online Active Learning

Next, we present the results of the online active learning experiments. We sequentially describe the results of three experiments: single-objective active learning aimed at improving ΔΔ*G_FlexddG_* alone, multi-objective optimization, and bispecific antibody exploration targeting both 5A12_Ang2 and 5A12_VEGF.

#### Single-Objective Sequence Sampling

Figure 6 shows the evolution of TopMean@40 and the Spearman correlation for ALLM-Ab and GPR(AbLang2) across different mutant generation methods and the existing AntBO.

**Figure 6:**
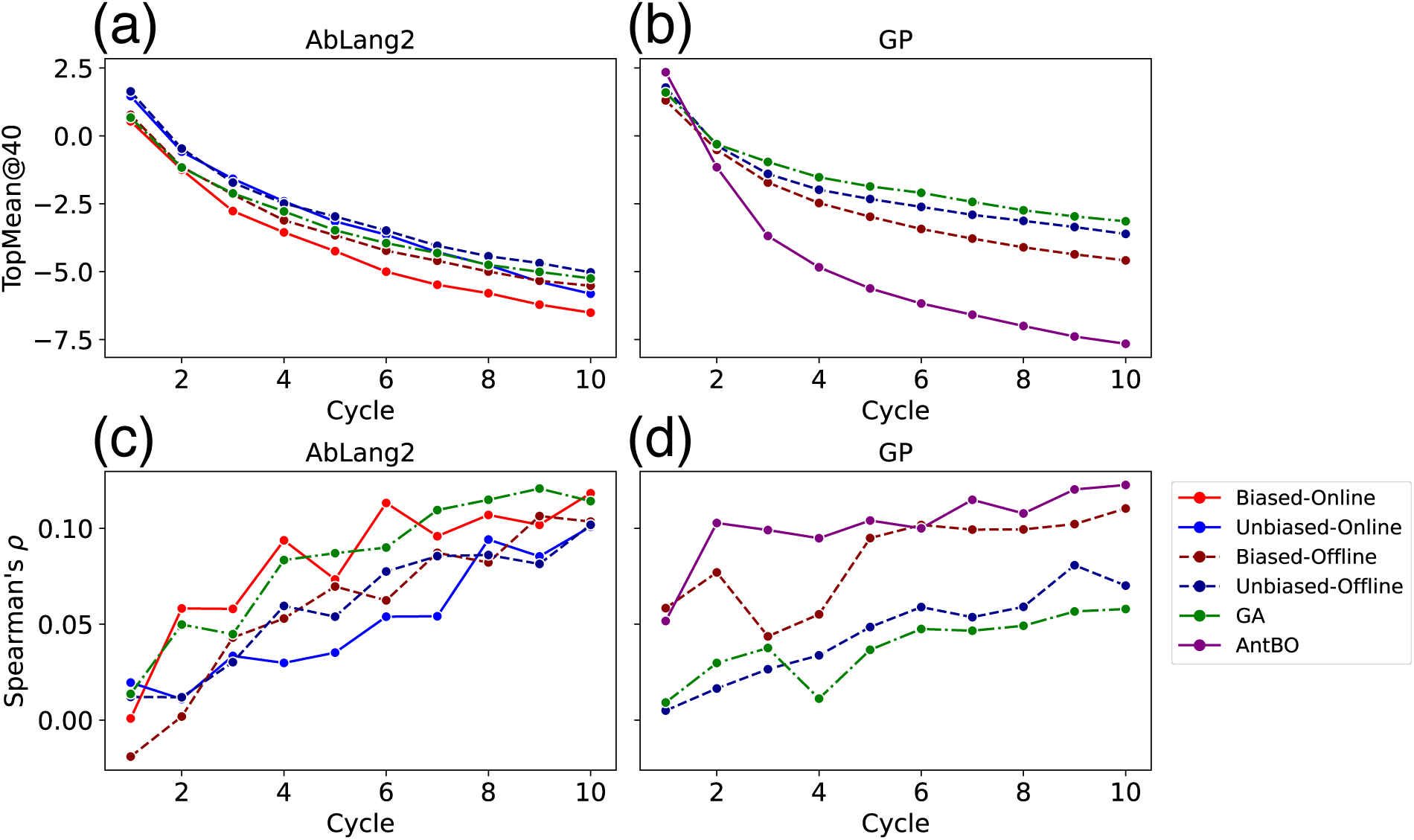
Evolution of TopMean@40 of ΔΔ*G_FlexddG_* and Spearman’s *ρ* for ALLM-Ab and GPR across five mutant generation methods. (a) TopMean@40 for ALLM-Ab, (b) TopMean@40 for GPR, (c) Spearman’s *ρ* for ALLM-Ab, (d) Spearman’s *ρ* for GPR.

First, regarding TopMean@40 values, the existing AntBO showed higher performance than ALLM-Ab and GPR(AbLang2), and no superiority of ALLM-Ab was observed. However, it should be noted that AntBO hardly considers antibody-like sequence characteristics, so there is a possibility that it falls into sequences that deviate from practical antibody space. This point will be verified in the next section. Then, TopMean@40 values indicate that ALLM-Ab was successful in discovering sequences with lower energy values, while for GPR(AbLang2), even the best performing case (Biased-Online) showed worse active learning performance than ALLM-Ab’s worst case (Unbiased-Offline). GPR(AbLang2) exhibited overall poor Spearman correlation, which explains why its TopMean@40 did not improve. Unlike the offline active learning using BindingGYM’s DMS data, the generated mutants are not restricted in the number of mutations from the wild-type sequence, making predictive accuracy for multi-point mutations crucial. This suggests that the loss of detailed amino acid residue information in ALLM-Ab’s latent space prevented GPR(AbLang2) from achieving sufficient predictive accuracy to effectively guide the active learning process.

When comparing GA-based generation to language model-based sampling, for ALLM-Ab the performance of GA and Biased-Offline were comparable. However, for GPR(AbLang2), the use of GA resulted in no improvement in Spearman correlation from the beginning, suggesting that promising mutants could not be discovered, likely due to poor extrapolation on out-of-distribution data by the GPR model. Note that the GA used here is a naïve repetition of independent GA runs for each cycle, resulting in extremely high computational cost. In contrast, sequence generation via language models, when combined with approximate fitness score for fast evaluation, offers little advantage to using GA given the associated cost.

Next, among the four language model-based sequence generation methods, applying bias using wild-type residues consistently resulted in higher Spearman correlation and a greater ability to discover sequences with low energy. In the absence of such bias, the vastness of the generated sequence space may lead the model to select out-of-distribution mutants.

Additionally, for ALLM-Ab, online sampling by generating sequences on the fly was more effective at discovering low-energy mutants. Overall, these results indicate that dynamically generating candidate sequences using a fine-tuned language model contributes to the discovery of sequences with lower energy values, and that constraining the search space using wild-type residue bias is important.

Next, Figure 7 shows the evolution of TopMean@40 of ΔΔ*G_FlexddG_* and Spearman correlation when using normal fitness scores versus approximation scores. Similar to the offline setting, it was confirmed that when using approximation scores, although Spearman correlation decreases, active learning performance equivalent to normal fitness scores can be obtained. As shown in Figure S1, using approximation scores can significantly reduce inference time.

**Figure 7:**
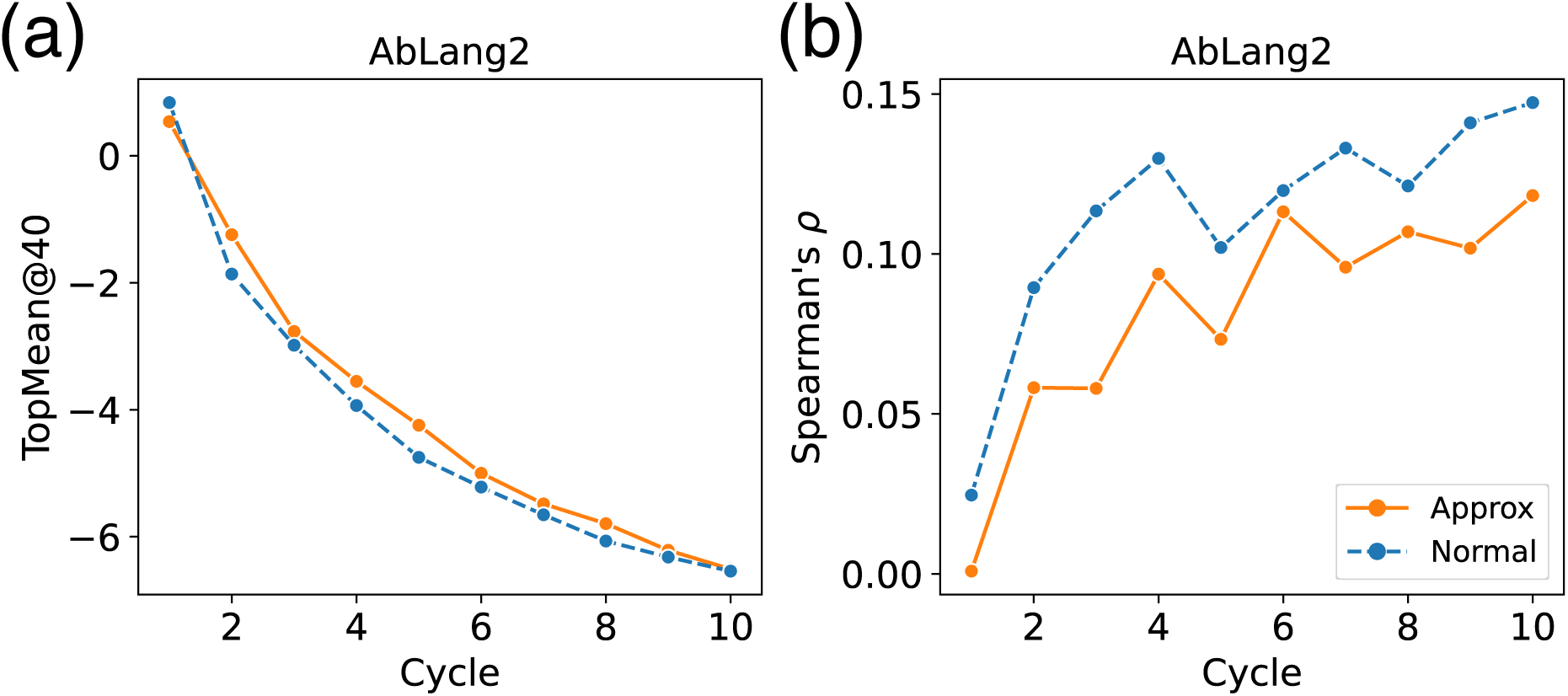
Evolution of TopMean@40 of ΔΔ*G_FlexddG_* and Spearman’s *ρ* when using fitness score (normal mode) versus approximation score (approx mode) in ALLM-Ab. (a) TopMean@40, (b) Spearman’s *ρ*.

#### Multi-Objective Optimization

Figure 8 shows scatter plots of the evolution of Flex ddG and developability metrics for mutants selected by each multi-objective optimization method. Also, Figure 9 shows the distribution of each objective variable for mutants ranked highly in the final cycle. For specific sequence logos, refer to Figures S2 through S9.

**Figure 8:**
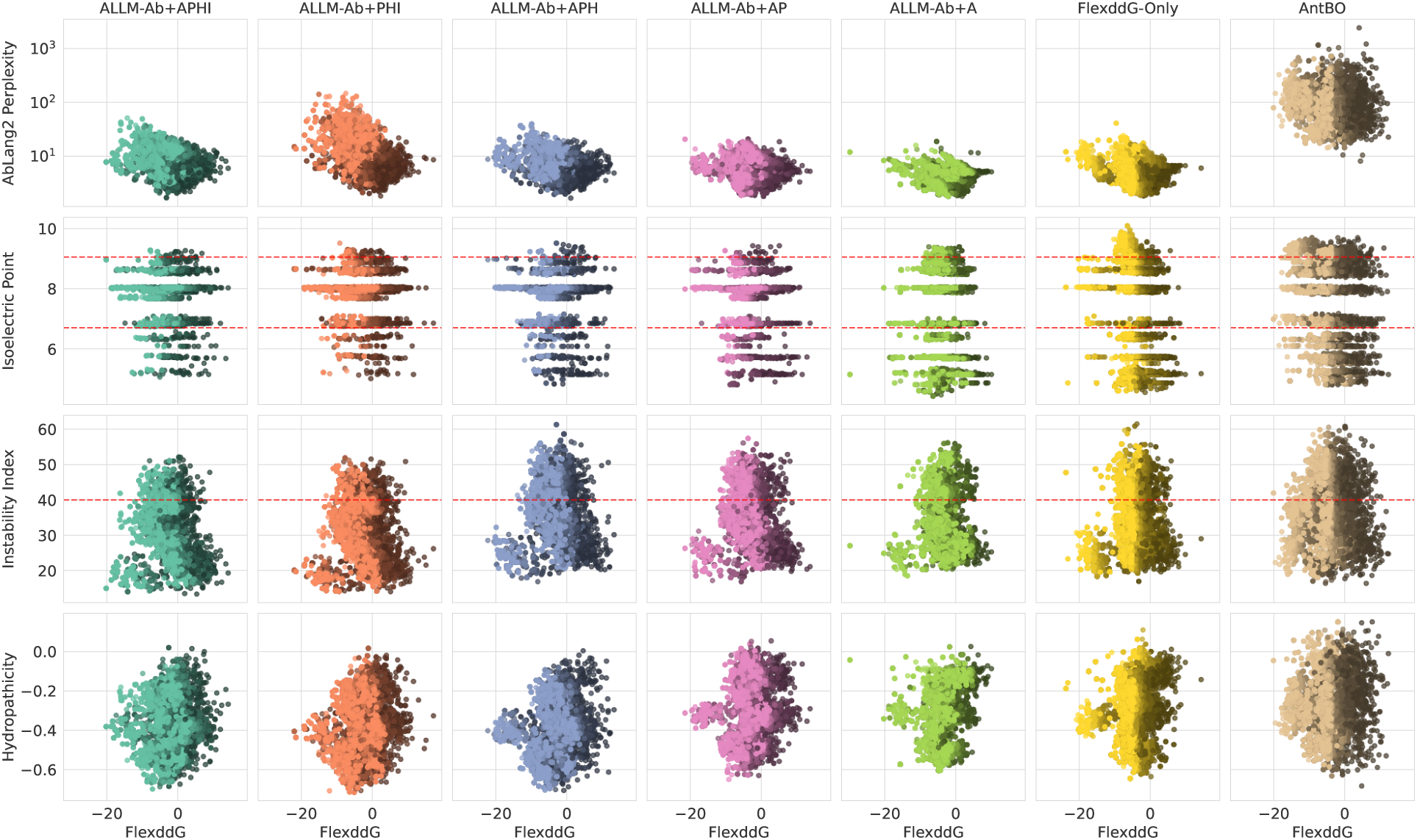
Scatter plots showing the evolution of Flex ddG and developability metrics for the top 40 mutants selected by each multi-objective optimization method in each cycle. Brighter colored points represent mutants that were ranked highly in later cycles.

**Figure 9:**
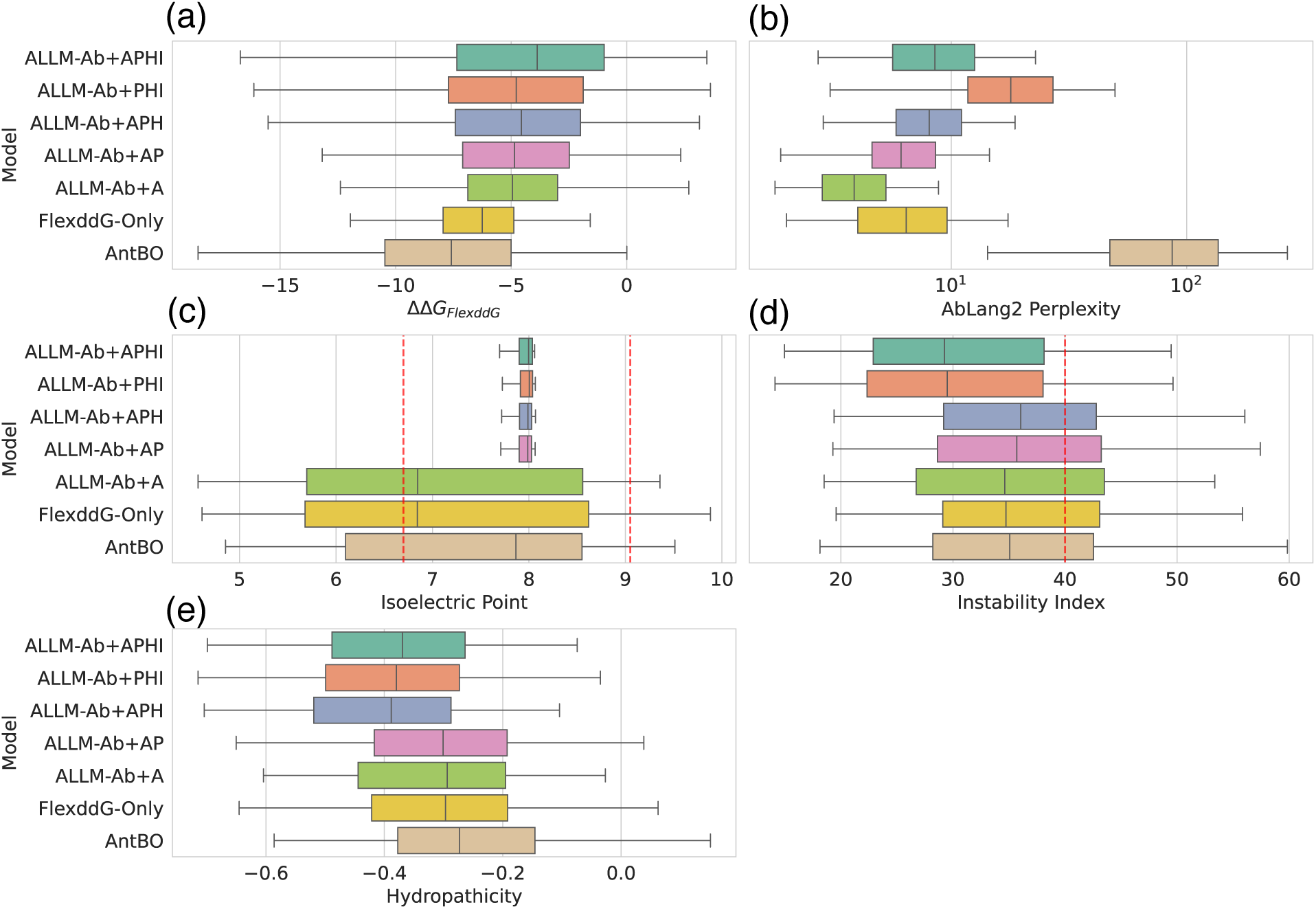
Box plots showing the distribution of objective variables for the top 40 mutants finally selected by each multi-objective optimization method. a) ΔΔ*G_FlexddG_*, b) AbLang2 perplexity score, c) Isoelectric point, d) Instability index, e) Hydropathicity.

First, as can be seen from Figure 9a, as the number of objective variables used for optimization increases, the Flex ddG values tend to decrease. This suggests a trade-off relationship between developability metrics and Flex ddG. AntBO selects mutants with the lowest Flex ddG values, but has the highest hydropathicity metric. Also, since it does not consider AbLang2 perplexity score, there is a tendency not to select antibody-like sequences. Indeed, looking at Figure S9, in many targets, selection is biased toward hydrophobic residues, and there is a high possibility of non-specifically lowering energy. Additionally, it is known that Arg/Lys at H94 and Asp at H101 are highly conserved in human antibody CDR-H3. ^73^ However, AntBO’s output sequences do not select these amino acids, resulting in significant deviation from standard CDR-H3. As ALLM-Ab+A, ALLM-Ab+AP, ALLM-Ab+APH,

ALLM-Ab+APHI are added with each optimization target, each developability metric improves. Although this is obvious since they are introduced as objective variables, it can be said that mutant exploration based on hypervolume maximization appropriately achieves multi-objective optimization. In ALLM-Ab+A, by keeping AbLang2 perplexity score low, antibody-specific sequence features appear as can be seen from Figure S7, but tyrosine is excessively selected in 1FBI and 1WEJ, etc. This is because AbLang2 recognizes CDR-H3 with many tyrosines as antibody-like sequences, and tyrosine can be introduced to lower Flex ddG depending on the target, so it is considered to be selected in a biased manner by these two factors. Such bias was improved by introducing the instability index as an objective variable.

Next, Figure 10 shows the distribution of antibody developability metrics for the top-1 mutants calculated for external evaluation. First, regarding PPC, only ALLM-Ab+APH shows significant deviation, but this is improved in ALLM-Ab+APHI through the introduction of the instability index. SFvCSP shows overall improvement through the introduction of isoelectric point. SAPpos shows high values in all methods that do not consider hydropathicity. For OASis, AntBO shows reduced human antibody-likeness, and ALLM-Ab+PHI shows decreased OASis, which corresponds to the AbLang2 perplexity score results in Figure 9b. Therefore, improving AbLang2 perplexity score was effective in considering human antibody-likeness. From the above results, it was confirmed that multi-objective optimization is important for antibody developability in external evaluation. However, introducing more developability also involves trade-offs with affinity, so it should be carefully considered in practical situations.

**Figure 10:**
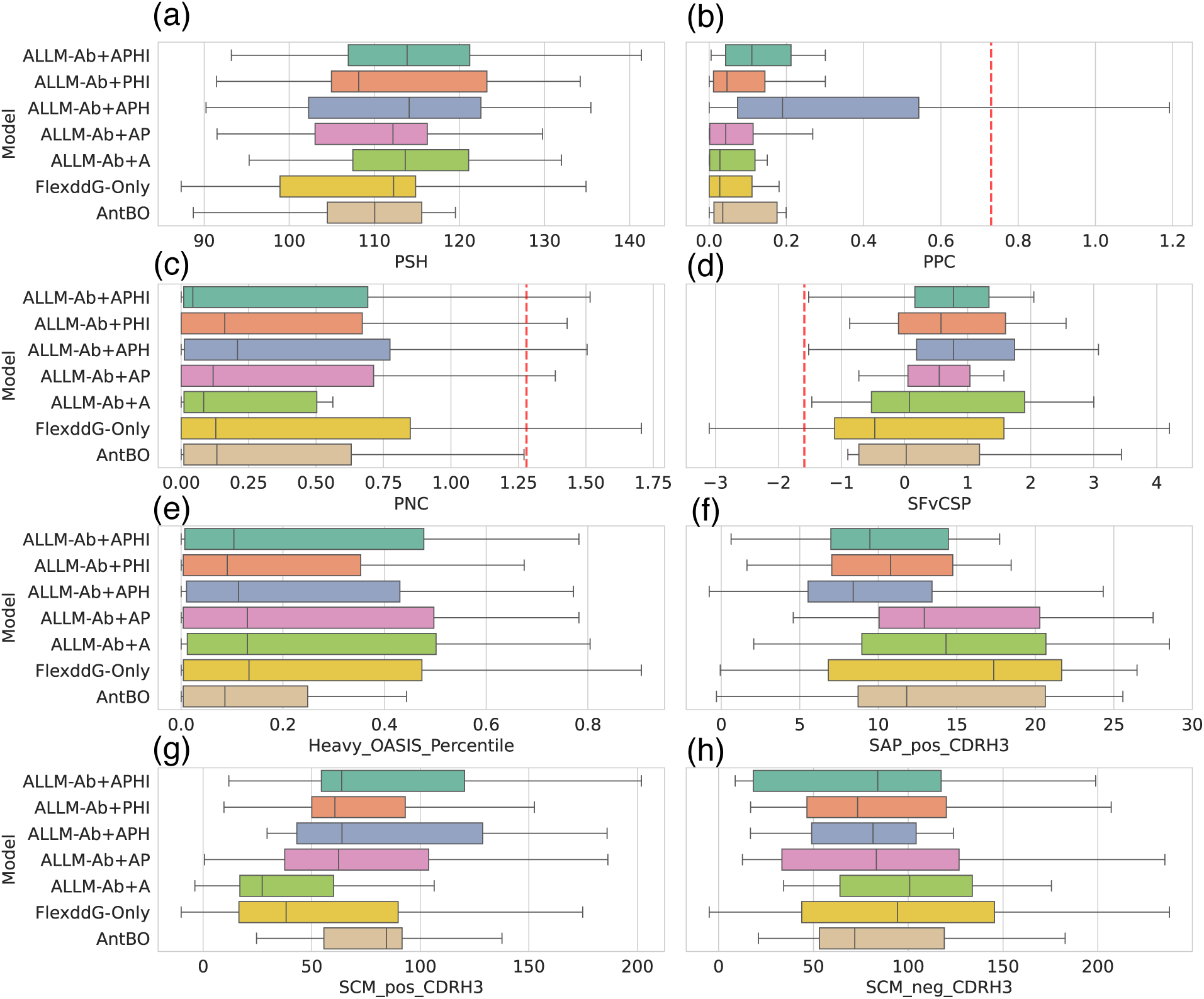
Box plots showing the distribution of antibody developability metrics for the top 1 mutant from each multi-objective optimization method. a) PSH, b) PPC, c) PNC, d) SFvCSP, e) Heavy OASis Percentile, f) `SAP_pos_CDR-H3`, g) `SCM_pos_CDR-H3`, h) `SCM_neg_CDR-H3`.

#### Dual Optimization for Ang2 and VEGF

Finally, we conducted dual optimization experiments for antibodies targeting both 5A12_Ang2 and 5A12_VEGF. Figure 11 shows the evolution of Flex ddG and developability metrics for mutants selected by FlexddG-Only and ALLM-Ab dual optimization. Also, Figure 12 shows the distribution of each objective variable for mutants ranked highly in the final cycle. Similar to the multi-objective optimization experiments, ALLM-Ab selected mutants with lower instability index and hydropathicity, preferentially selecting mutants with favorable characteristics across all considered properties. In contrast, ΔΔ*G_FlexddG_* for Ang2 and VEGF tends to be inferior to FlexddG-Only. This suggests a stronger trade-off between improving binding affinity and maintaining antibody sequence validity in the dual optimization setting. Also, FlexddG-Only is considered to non-specifically lower energy for both antibodies by biasing toward mutants with many hydrophobic residues.

**Figure 11:**
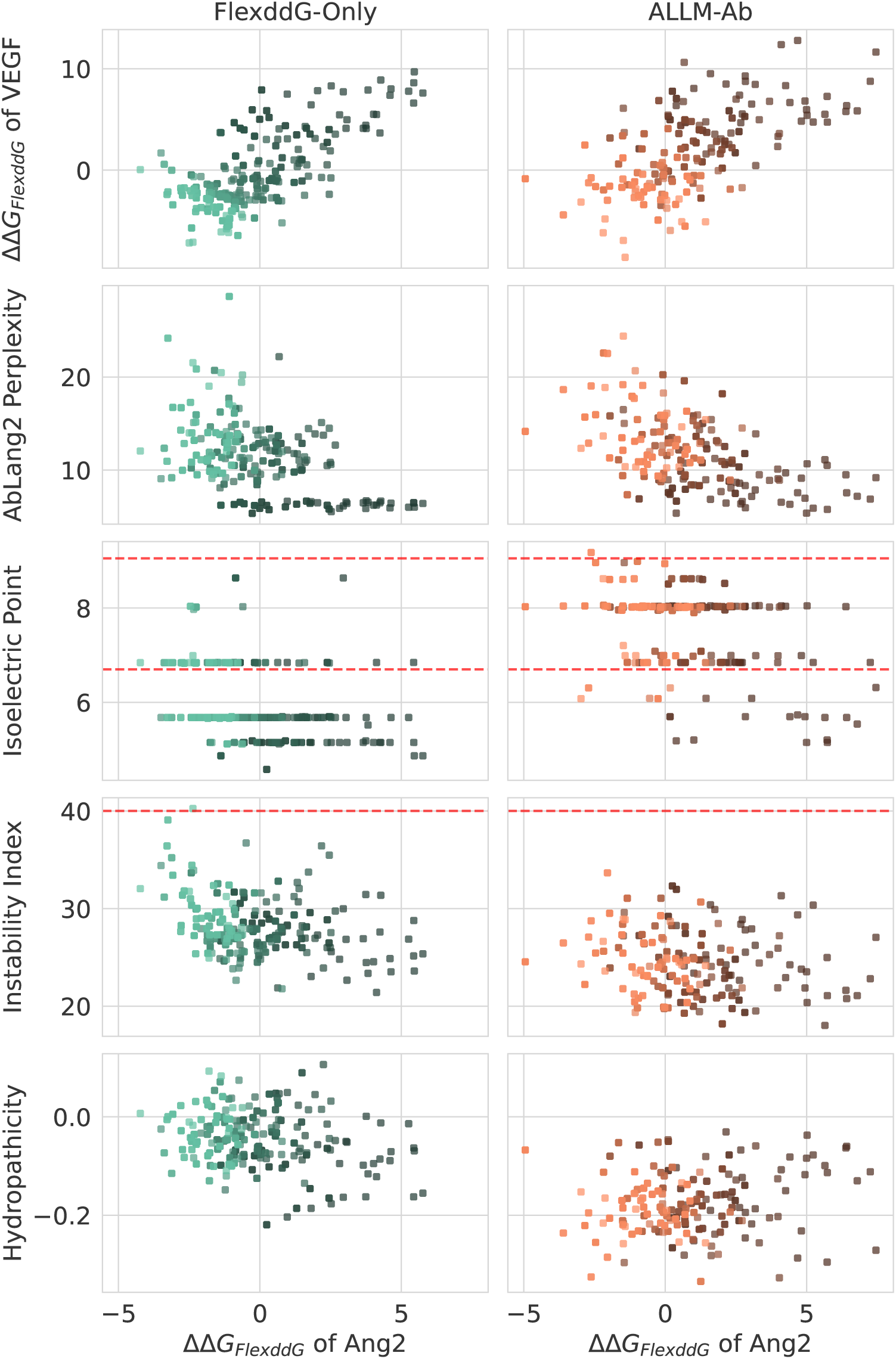
Scatter plots showing the evolution of Flex ddG and developability metrics for the top 40 mutants selected by dual optimization in each cycle. a) FlexddG-Only, b) ALLM-Ab. Brighter colored points represent mutants that were ranked highly in later cycles.

**Figure 12:**
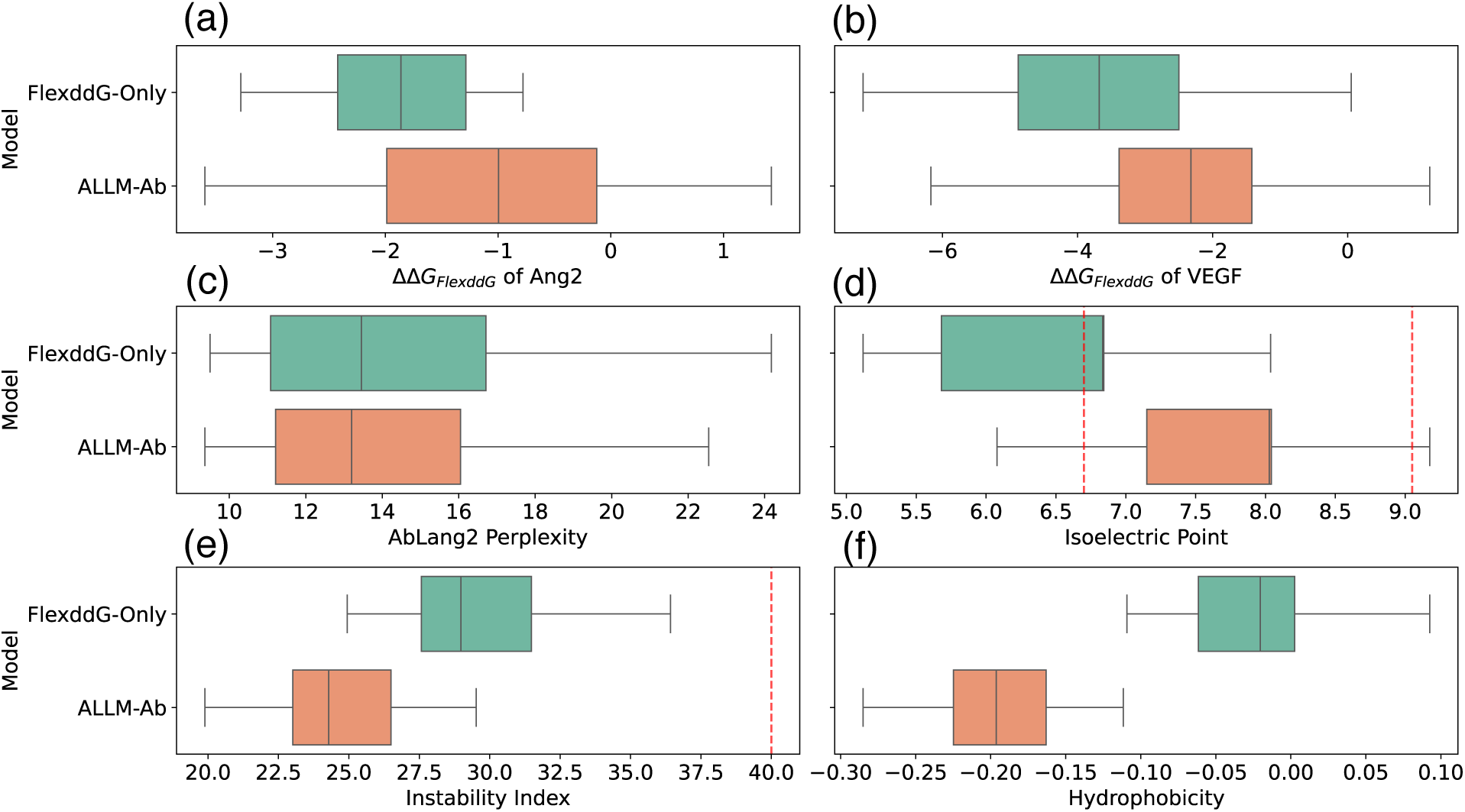
Box plots showing the distribution of objective variables for the top 40 mutants finally selected by dual optimization. a) ΔΔ*G_FlexddG_* for Ang2, b) ΔΔ*G_FlexddG_* for VEGF, c) AbLang2 perplexity score, d) Isoelectric point, e) Instability index, f) Hydropathicity.

Also, Figure 13 shows sequence logos of the top 40 mutants selected by dual optimization. Similar to the multi-objective optimization experiments, FlexddG-Only tends to excessively select bulky hydrophobic residues such as phenylalanine, and there is a high possibility of non-specifically lowering energy. ALLM-Ab succeeded in suppressing this, but there is still a bias toward hydrophobic residues. This suggests that in dual optimization, it is difficult to optimize energy while suppressing hydropathicity.

**Figure 13:**
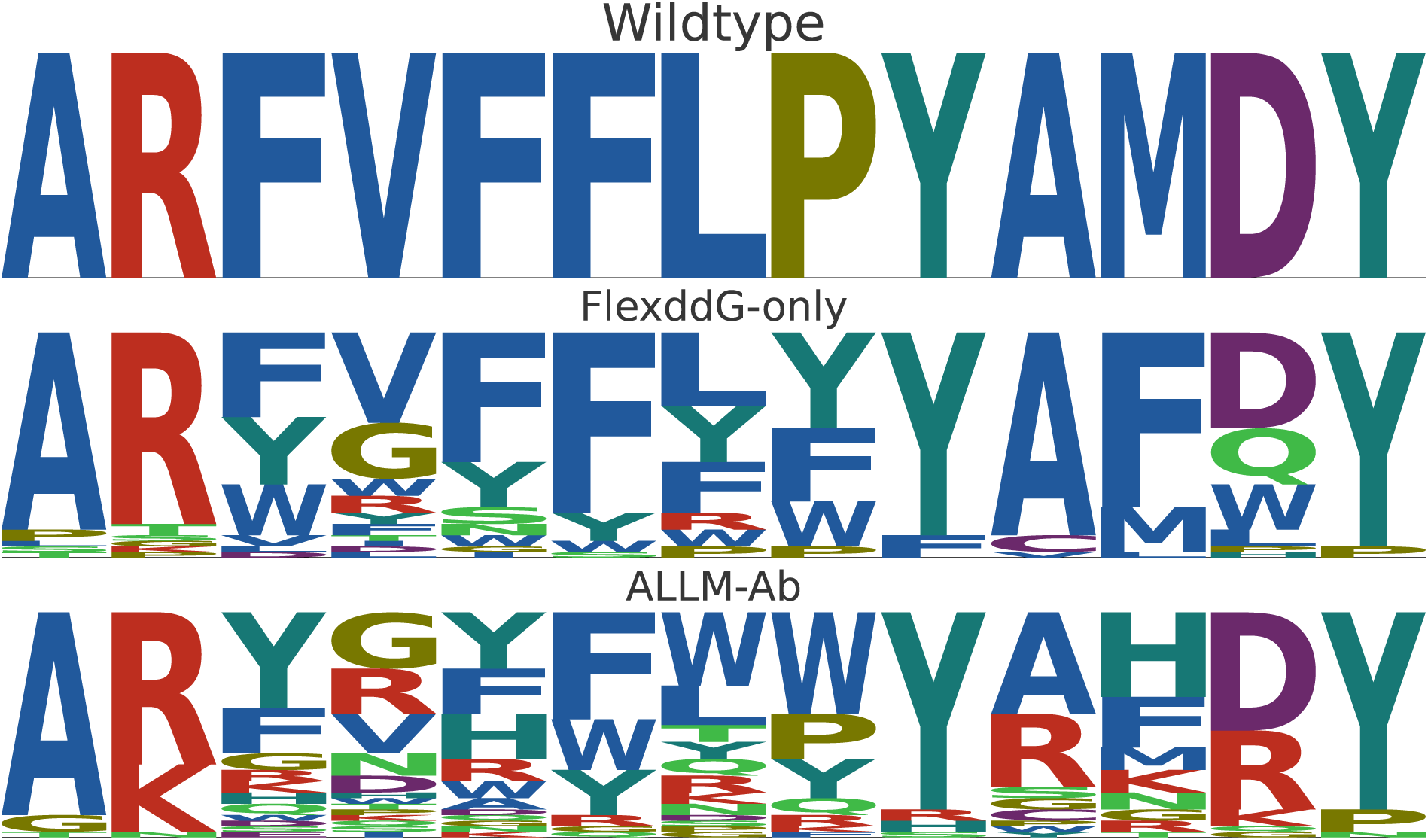
Sequence logos of the top 40 mutants selected by dual optimization. The first row shows the wild-type sequence, the second row shows the results for FlexddG-Only, and the third row shows the results for ALLM-Ab.

Finally, Figure 14 shows examples of structures optimized with Flex ddG for sequences selected by dual optimization. For Ang2, both the wild-type (Figure 14a) and optimized mutants (Figure 14b) maintain hydrogen bonds with cysteine and methionine of Ang2, preserving important interactions. For VEGF, while the wild-type (Figure 14c) maintains hydrogen bonds with histidine and glutamine of VEGF, the optimized mutant (Figure 14d) additionally includes hydrogen bonds between the 36th lysine of VEGF and tyrosine of CDR-H3, as well as cation-*π* interactions with tryptophan of CDR-H3. Thus, it was confirmed that dual optimization by ALLM-Ab achieved higher binding affinity by appropriately adding new interactions while maintaining important interactions specific to the two targets.

**Figure 14:**
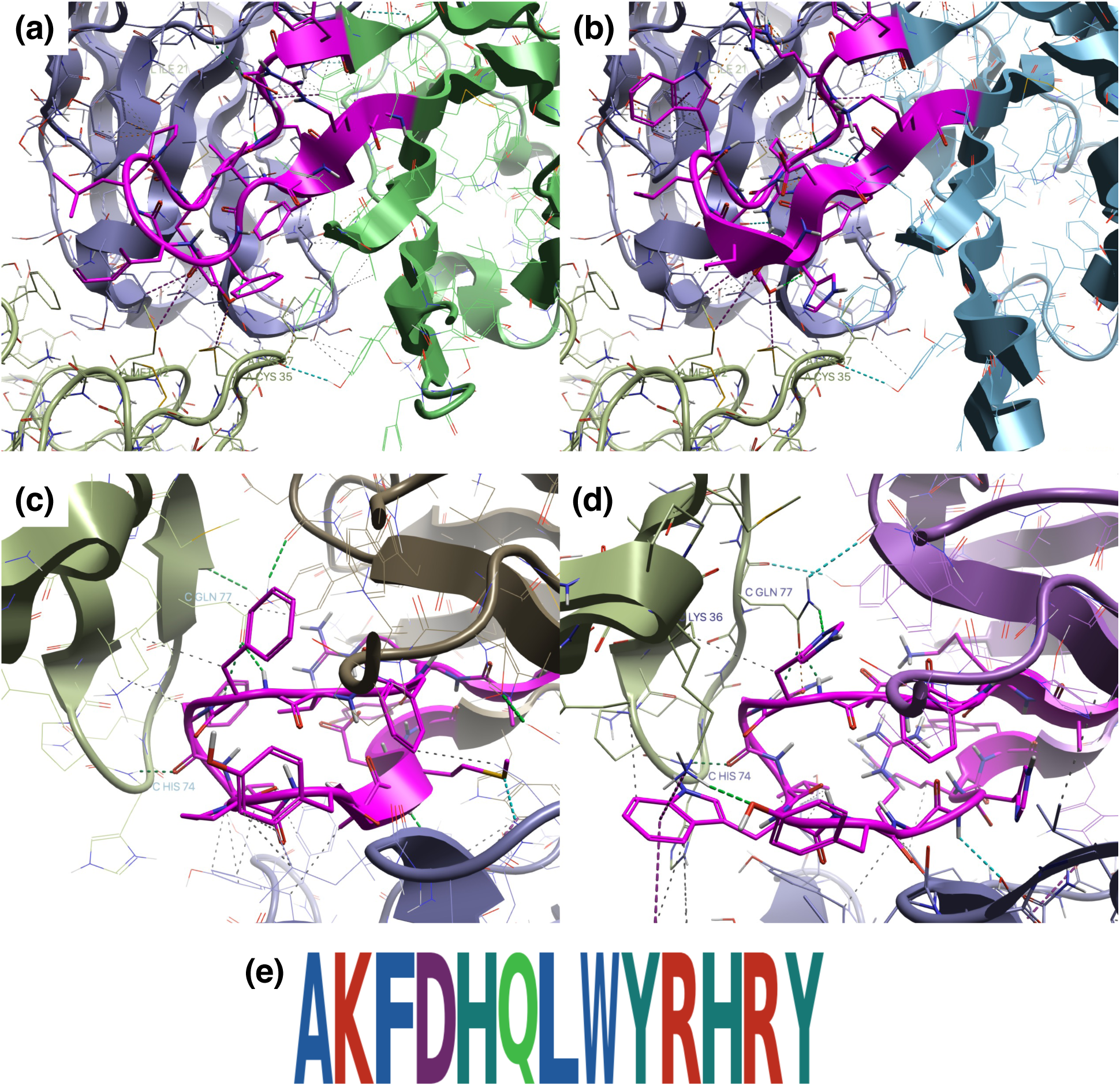
a) Wild-type complex structure for Ang2, b) Example structure selected by dual optimization for Ang2, c) Wild-type complex structure for VEGF, d) Example structure selected by dual optimization for VEGF, with pink representing CDR-H3. e) CDR-H3 sequence in this example.

## Conclusions

In this study, we proposed ALLM-Ab, an active learning approach for antibody sequence optimization using protein language models. ALLM-Ab integrates three key components: (1) parameter-efficient fine-tuning with learning-to-rank for sequence scoring, (2) efficient sequence sampling via the fine-tuned model, and (3) multi-objective optimization considering antibody-likeness and developability. We evaluated ALLM-Ab in two scenarios: offline active learning using BindingGYM data without sequence generation and online active learning aimed at optimizing Flex ddG energy values. In the offline active learning experiments, although a conventional latent space-based Gaussian process regression (GPR) approach exhibited higher predictive performance, it struggled in the online setting due to poor extrapolation to out-of-distribution data. In contrast, ALLM-Ab based on fine-tuned language models for sequence sampling was able to directly reflect the model’s preferences, thereby achieving more efficient optimization compared to genetic algorithm-based approaches. Moreover, we showed that approximation scores lead to a significant reduction in computation time with only a modest decrease in performance for active learning.

In the online active learning experiments, incorporating multiple developability metrics as additional objectives enabled us to maintain high-developability antibody sequence features while still discovering mutants with high binding affinity. The existing state-of-the-art method AntBO was excellent at discovering mutants with high binding affinity, but tended to excessively select hydrophobic residues to easily gain energy scores, indicating that consideration of developability metrics is important. In the dual optimization experiments for bispecific antibodies targeting 5A12_Ang2 and 5A12_VEGF, a more pronounced trade-off between improving binding affinity and preserving sequence validity was observed compared to the single-target optimization case. Nevertheless, our proposed approach is applicable even in such complex optimization scenarios.

There are several important limitations to this study. First, the correlation between Flex ddG energy and actual binding affinity is limited. However, the proposed active learning approach is not limited to Flex ddG, which was used in this study as a surrogate for evaluation. It can be generalized to other methods, such as low-throughput free energy perturbation (FEP) calculations^25^ or experimental measures like ELISA assay.^23^ Nevertheless, because FEP calculations can only evaluate a limited number of mutations at once, active learning strategies that take mutation cost into account are required. Furthermore, important limitations of this study include the lack of verification for optimization beyond CDR-H3 and the inability to change the length of CDR-H3 during the optimization process. Additionally, since pplscore alone was insufficient to achieve developability, it was necessary to introduce multiple developability metrics. By learning therapeutic antibody-like characteristics through antibody language models, it might become possible to perform optimization limited to more appropriate antibody sequence spaces. Future work should include experimental validation of the proposed approach and its application to scenarios with limited mutations (e.g., relative FEP calculations). This research presents a novel active learning approach that leverages the characteristics of protein language models, with the potential to accelerate future antibody development research and reduce the increasing development costs of therapeutic antibodies.

## Data and Software Availability

The ALLM-Ab code and the dataset used in this study are available at https://github.com/ohuelab/ALLM-Ab.

## Supporting Information Available

Supporting information is available free of charge.

- SI.pdf: Additional figures about detailed experimental results.

## Acknowledgement

The authors acknowledge the use of Claude-4-Sonnet for the initial translation of this manuscript from Japanese to English. The authors subsequently verified the scientific accuracy and appropriateness of technical terminology, and made final revisions to ensure the quality of the manuscript.

