## Supporting Information for "ALLM-Ab: Active Learning-Driven Antibody Optimization Using Fine-tuned Protein Language Models"

Table S1: Configuration of LoRA target layers for each model.

| Model | Target Modules |
|---|---|
| ProteinMPNN | `W1`, `W2`, `W3`, `W11`, `W12`, `W13`, `W_in`, `W_out` |
| ESM2 | `q_proj`, `k_proj`, `v_proj`, `out_proj`, `lm_head.dense` |
| AbLang2 | `q_proj`, `k_proj`, `v_proj`, `out_proj`, `intermediate_layer.0`, `intermediate_layer.2` |

Table S2: Comparison of average time for sampling 100,000 sequences with and without applying approximation score.

| Method | Execution Time (min) |
|---|---|
| Normal | 17.0 |
| Approx | 1.12 |

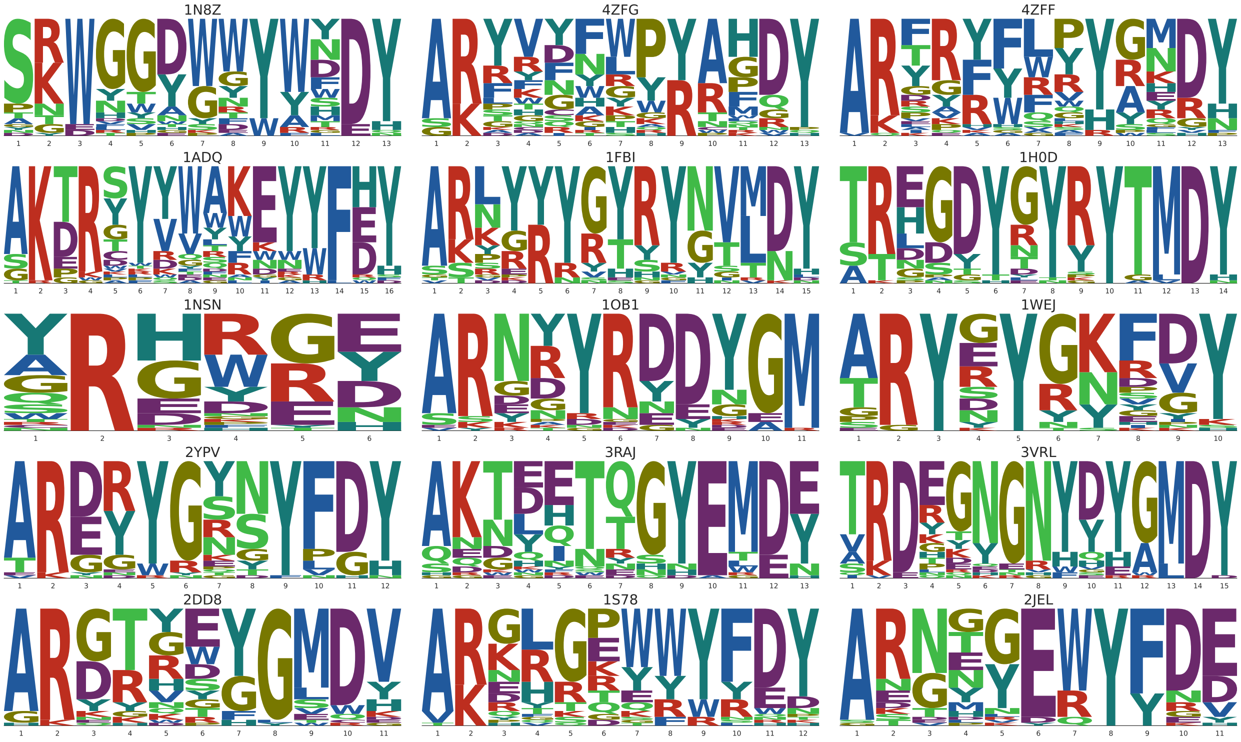


Figure S1: Sequence logos of the top 40 mutants selected by ALLM-Ab+APHI for 15 targets.

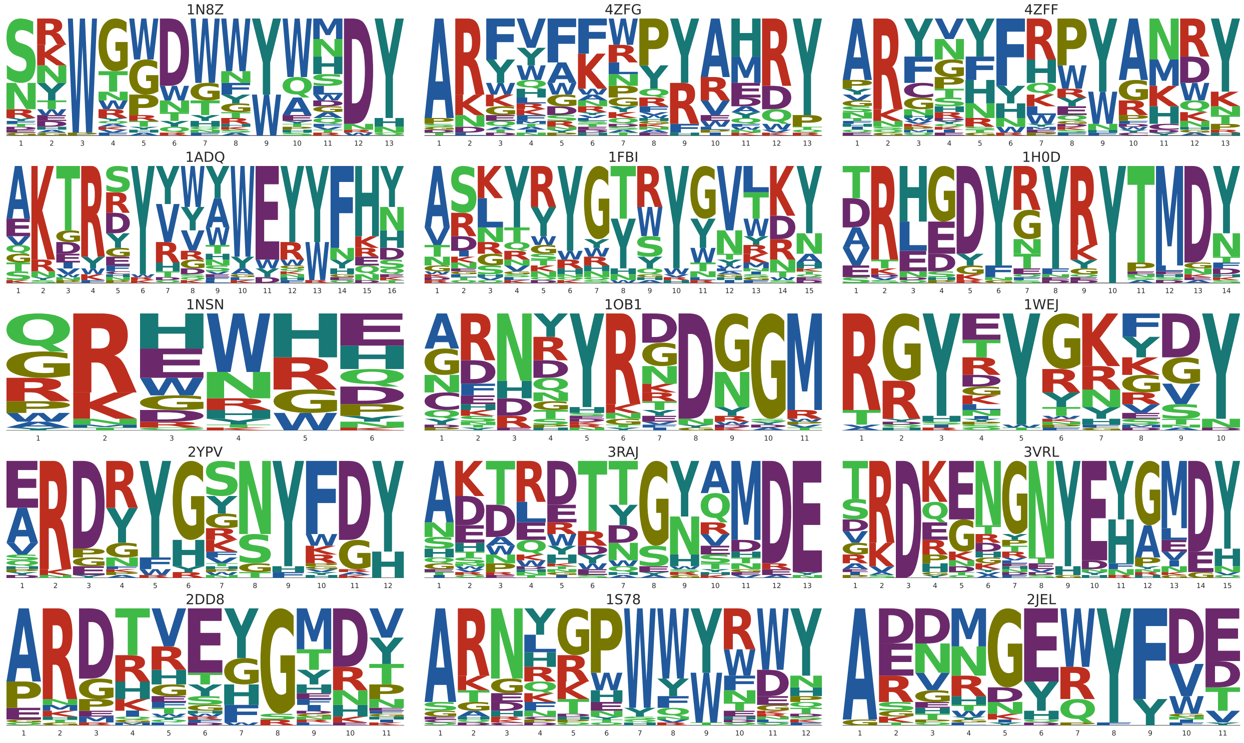


Figure S2: Sequence logos of the top 40 mutants selected by ALLM-Ab+PHI for 15 targets.

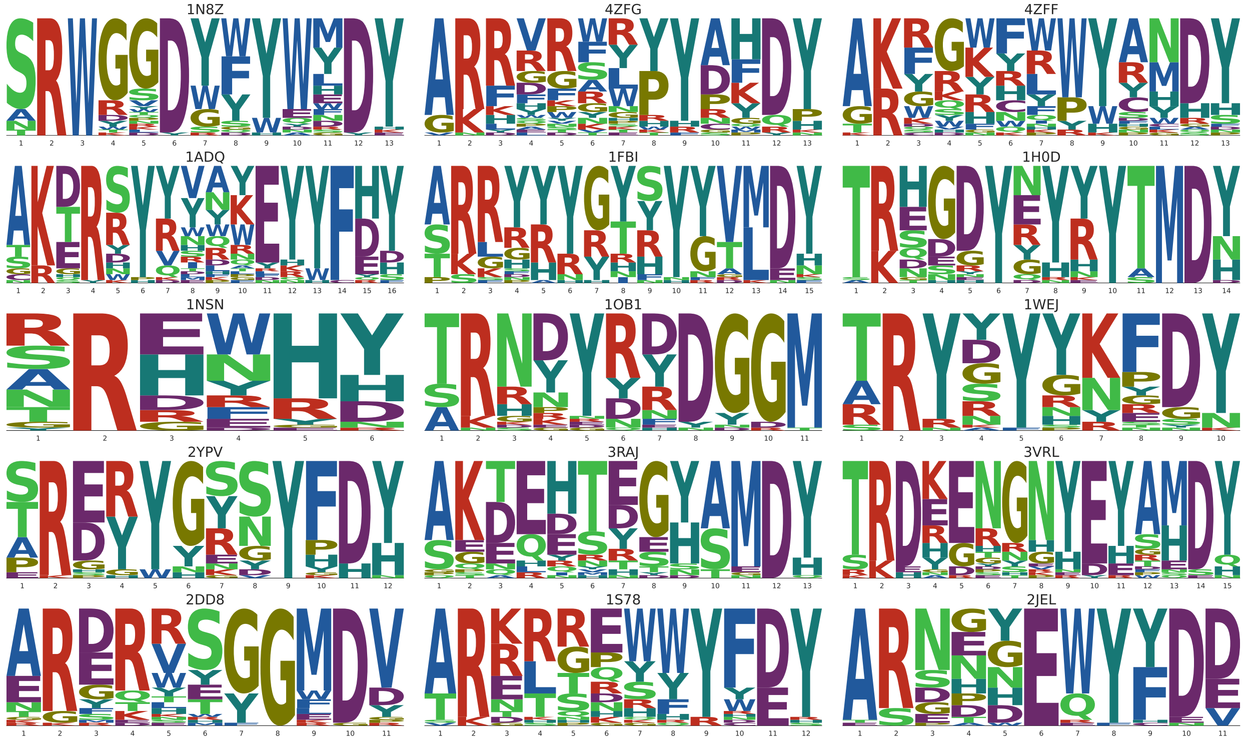


Figure S3: Sequence logos of the top 40 mutants selected by ALLM-Ab+APH for 15 targets.

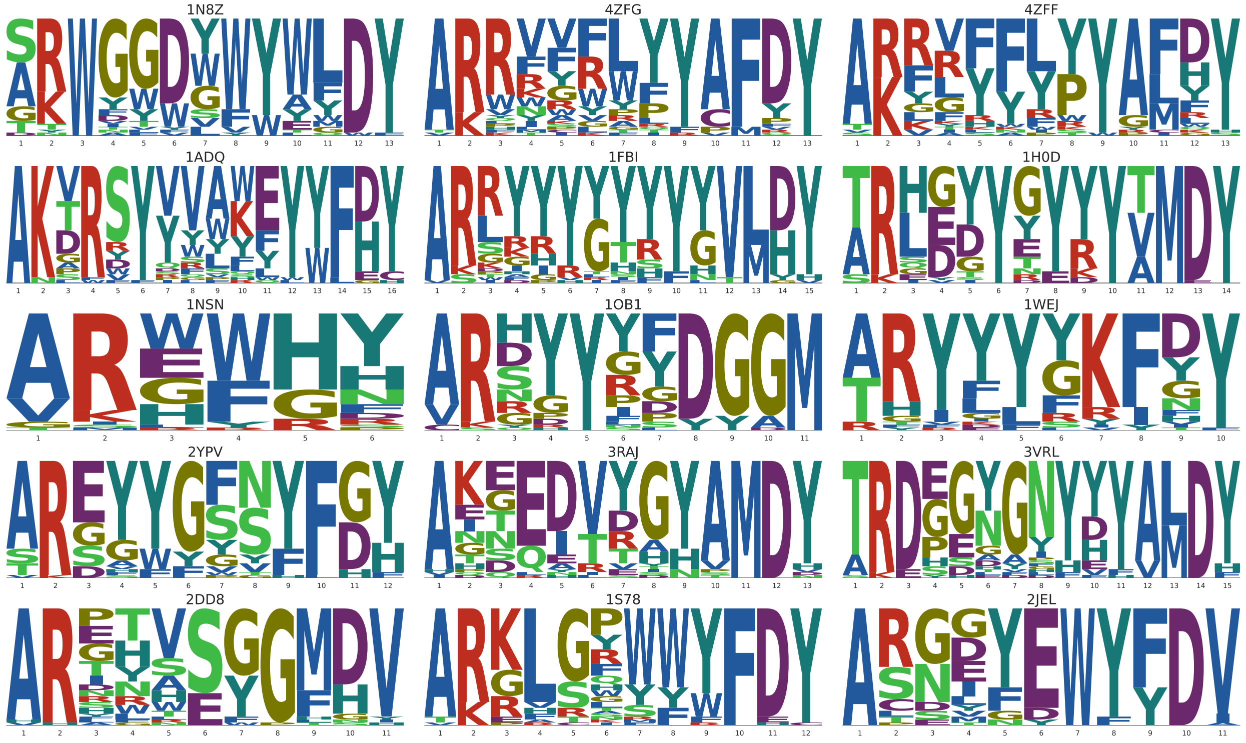

Figure S4: Sequence logos of the top 40 mutants selected by ALLM-Ab+AP for 15 targets.

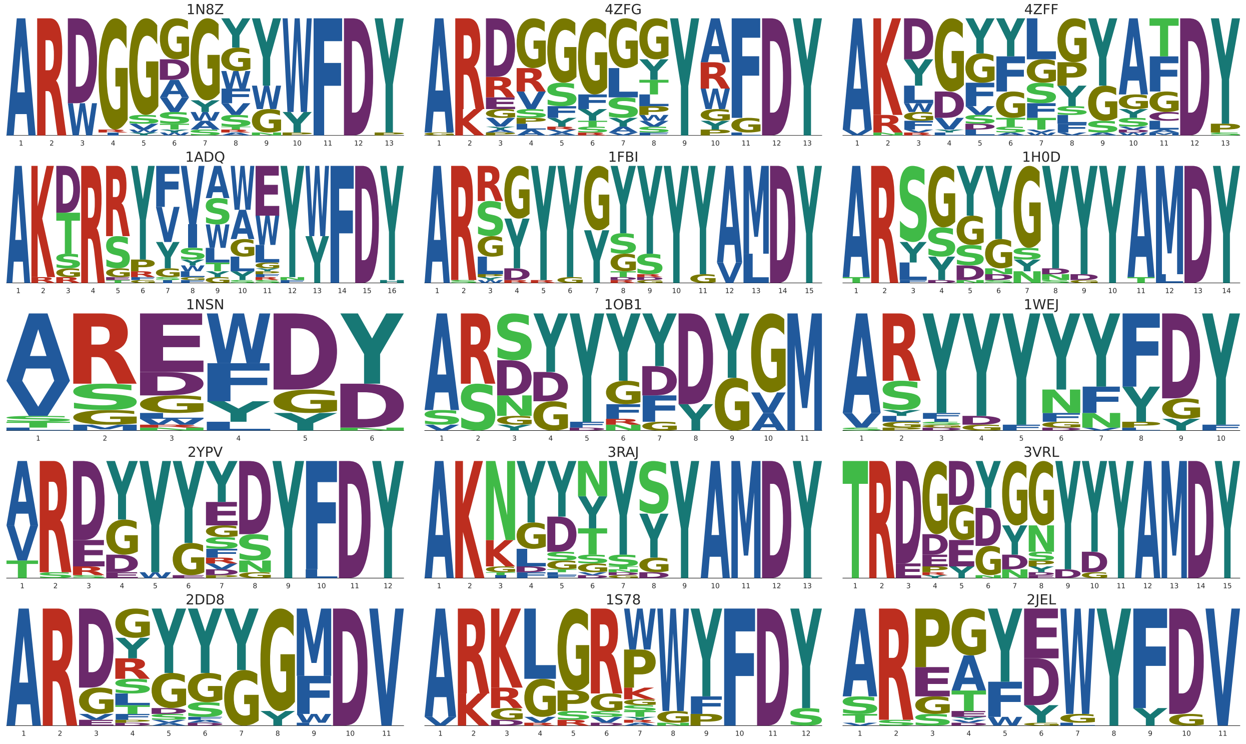

Figure S5: Sequence logos of the top 40 mutants selected by ALLM-Ab+A for 15 targets.

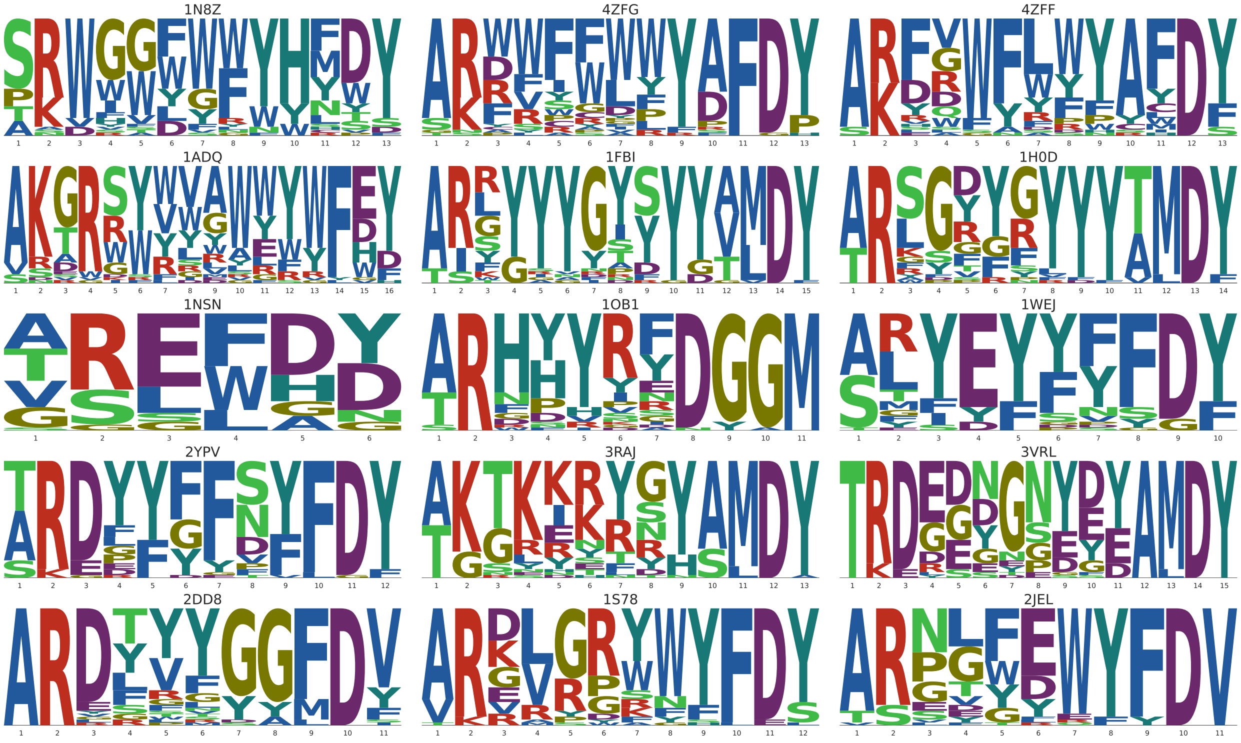


Figure S6: Sequence logos of the top 40 mutants selected by FlexddG-Only for 15 targets.

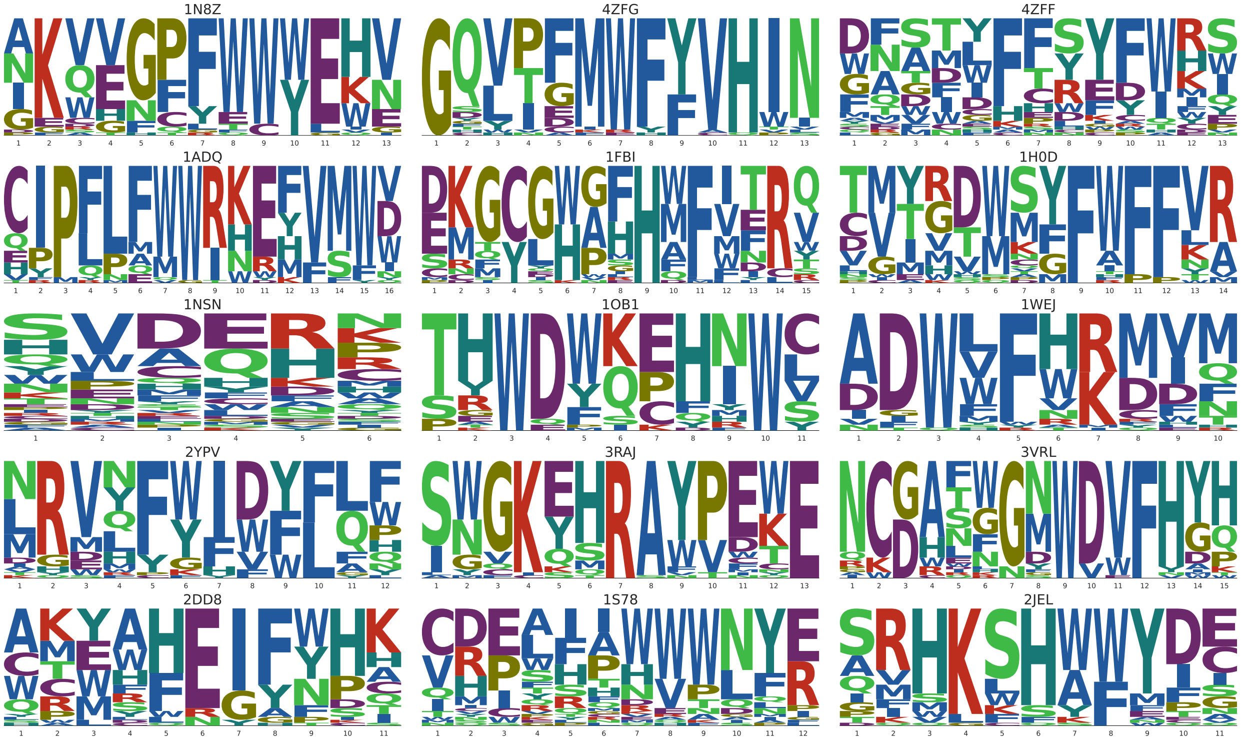


Figure S7: Sequence logos of the top 40 mutants selected by AntBO for 15 targets.
